# Transcriptional perturbation of LINE-1 elements reveals their *cis*-regulatory potential

**DOI:** 10.1101/2024.02.20.581275

**Authors:** Yuvia A Pérez-Rico, Aurélie Bousard, Lenka Henao Misikova, Eskeatnaf Mulugeta, Sérgio F de Almeida, Alysson R Muotri, Edith Heard, Anne-Valerie Gendrel

## Abstract

Long interspersed element-1 (LINE-1 or L1) retrotransposons constitute the largest transposable element (TE) family in mammalian genomes and contribute prominently to inter- and intra-individual genetic variation. Although most L1 elements are inactive, some evolutionary younger elements remain intact and genetically competent for transcription and occasionally retrotransposition. Despite being generally more abundant in gene-poor regions, intact or full-length L1s (FL-L1) are also enriched around specific classes of genes and on the eutherian X chromosome. How proximal FL-L1 may affect nearby gene expression remains unclear. In this study, we aim to examine this in a systematic manner using engineered mouse embryonic stem cells (ESCs) where the expression of one representative active L1 subfamily is specifically perturbed. We found that ∼1,024 genes are misregulated following FL-L1 activation and to a lesser extent (∼81 genes), following their repression. In most cases (68%), misexpressed genes contain an intronic FL-L1 or lie near a FL-L1 (<260 kb). Gene ontology analysis shows that upon L1 activation, up-regulated genes are enriched for neuronal function-related terms, suggesting that some L1 elements may have evolved to control neuronal gene networks. These results illustrate the *cis*-regulatory impact of FL-L1 elements and suggest a broader role for L1s than originally anticipated.

## Introduction

Transposable elements (TEs) constitute nearly half of mammalian genomes. These sequences have long been considered purely parasitic elements self-propagating in the host genome. However, it is now widely accepted that TEs are major drivers of genome evolution and genetic diversity, capable of expanding gene regulatory networks (Chuong et al. 2017), as originally proposed by Barbara McClintock (McClintock 1956). TEs can be broadly classified into two classes: DNA transposons, which are mostly extinct in mammals, and retrotransposons, which transpose using a copy-paste mechanism and have massively contributed to the expansion of mammalian genome size. Among retrotransposons, long interspersed nuclear element-1 (LINE-1 or L1) constitute the most abundant TE family, representing ∼20% of the human and mouse genomes and the only known autonomous class of TE mobile in humans (Hermant and Torres-Padilla 2021).

The majority of L1 elements in mammalian genomes have been inactivated by the accumulation of mutations or truncations at their 5’ end. However, a small fraction of non-truncated L1 elements, known as full-length L1s (FL-L1s), still possess intact ORF1 and ORF2 sequences and retain the ability to retrotranspose. Additionally, a larger proportion of non-intact FL-L1s are characterised by disrupted ORF sequences, therefore losing the ability to retrotranspose autonomously, and populate mammalian genomes (Penzkofer et al. 2017). Many of these non-intact FL-L1s have retained a functional 5’UTR region and still have the potential to be transcriptionally active. As such, they may represent a hidden source of regulatory sequences capable of contributing massively to gene regulation and transcriptome diversity (Fueyo et al. 2022). Transcriptionally competent FL-L1s are stringently controlled both at the chromatin level, by epigenetic mechanisms and transcription factors, and at the post-transcriptional level. These mechanisms appear to act in a sequence-specific manner and depend on L1 evolutionary age or the cell type in which they operate (Bulut-Karslioglu et al. 2014; Castro-Diaz et al. 2014; Zoch et al. 2020). In the human genome, only ∼150 L1 copies are still capable of retrotransposition, all of them belonging to the young, human-specific L1Hs subfamily, while thousands of L1 copies remain transcriptionally competent (Brouha et al. 2003; Penzkofer et al. 2017). In the mouse genome, three young L1 subfamilies, called L1MdA, L1MdTf, and L1MdGf, are currently active. They differ mostly by their 5’UTR sequences containing the promoter and encompass thousands of transcriptionally competent elements (Sookdeo et al. 2013; Jachowicz and Torres-Padilla 2015; Penzkofer et al. 2017).

Uncontrolled activity of FL-L1s can have harmful consequences including DNA damage, genomic instability, gene disruptions and hindrance of the transcriptional coordination of gene networks. For that reason, FL-L1s are strongly silenced in somatic cells. Nevertheless, it has been observed that the activity of FL-L1s is enhanced upon aging and during malignancies (Payer and Burns 2019). Moreover, specific contexts have revealed distinct transcriptional dynamics of FL-L1s. For instance, significant transcriptional activity of FL-L1s has been observed in embryonic stem cells (ESCs) and during preimplantation development (Jachowicz et al. 2017; Richardson et al. 2017), or in neuronal progenitor cells (NPCs) and neurons in the hippocampus (Richardson et al. 2014; Upton et al. 2015). These observations raised the possibility of an actual functional role conferred to FL-L1 activity, for example, during reprogramming following fertilization (Jachowicz et al. 2017) or in neuronal somatic mosaicism and plasticity, either coordinating regulatory gene networks or creating genomic diversity (Richardson et al. 2014). Additionally, a subset of FL-L1s are transiently expressed from the X chromosome during X-chromosome inactivation, rather than being silenced, which could potentially facilitate gene silencing in specific regions of the chromosome (Chow et al. 2010; Loda et al. 2017).

The recent expansion of genome editing technologies and computational tools have accelerated our access to TEs and the possibility to devise approaches for direct and systematic interrogation of their potential functions (Goerner-Potvin and Bourque 2018; Fueyo et al. 2022). In recent years, various strategies have been developed to manipulate the expression of different classes of TEs, usually during early embryonic stages or in embryonic cells, contexts that are more permissive for their activity (Fueyo et al. 2022). For instance, a nuclease dead-Cas9 (dCas9) fused to effector domains was used to perturb the expression of LTR5Hs, a class of human endogenous retrovirus (ERVs), to show that these elements can act as early embryonic enhancers (Fuentes et al. 2018). TALEs (transcription activator-like effector) were used for transcriptional manipulation of endogenous L1 elements during mouse preimplantation development to demonstrate that L1 expression is critical for chromatin accessibility and developmental progression (Jachowicz et al. 2017). Finally, antisense oligonucleotides (ASOs) were used to target L1 RNA for degradation in mouse ES cells, which resulted in decreased self-renewal and proliferation, and demonstrated that L1 RNA appears to act as a nuclear RNA scaffold, regulating the expression of two-cell stage embryo genes (Percharde et al. 2018).

Given their abundance in mammalian genomes, TEs and their remnants are frequently found near or within genes. In certain situations, they have been co-opted as functional cis-regulatory elements. This phenomenon has been well documented on a genome-wide scale, for certain ERV families and SVAs elements in mouse or human cells (Hermant and Torres-Padilla 2021; Barnada et al. 2022; Fueyo et al. 2022). A recent study revealed that a significant portion of L1 repeats in mouse and human genomes are located within a short distance from genes (Lu et al. 2020). However, the fraction of FL-L1 elements that may contribute to gene expression regulation is unclear. In this study, we sought to investigate this further by studying the extent to which FL-L1 retrotransposons may act as *cis*-regulators and shape gene regulatory networks. To answer these questions and considering the elevated transcriptional activity of L1 elements in mouse ESCs (Martens et al. 2005), we used this model to engineer cell lines to perturb the expression of one representative young L1 subfamily in the mouse genome, L1MdTf. We report here the genome-wide specific repression and activation of targeted FL-L1s using zinc fingers-KRAB and dCas9-VPR systems. Hundreds of protein-coding genes were found to be misregulated following L1 perturbation, and these genes frequently contain an intronic L1 or lie near a L1 element targeted by the engineered effectors. Interestingly, we found that upregulated genes following L1 activation are enriched for neuronal function-related terms, in line with the observation that the activity of L1 elements is unleashed in neural progenitor cells and neurons (Richardson et al. 2014; Bodea et al. 2018).

## Results

### Engineered repressor and activator to manipulate L1MdTf expression

To investigate the impact of *cis*-regulation mediated by FL-L1 elements, we focused on the representative mouse young L1MdTf subfamily, which comprise an estimated number of 1,722 non-exonic non-intact FL elements (commonly defined as >= 6 kb) in RepeatMasker annotations (2021-04-08 version). To achieve genome-wide repression of L1MdTf elements, we took advantage of a fusion repressor composed of an array of 6 zinc-fingers (ZF) engineered to bind a 20 bp region within the 5’UTR monomeric consensus sequence of L1MdTf (Naas et al. 1998), fused to the KRAB repressor domain (called LKF) (Fig.1A, Fig.S1A). As a control, we used an analogous recombinant protein, carrying 4 amino-acid substitutions to alanine within each ZF, rendering them unable to bind DNA (called AKF). *In silico* analysis of the occurrence of the 20 bp region bound by each ZF predicted targeting of 1,064 out of 1,722 FL-L1MdTf, which comprise 93-95% of L1MdTf_I and L1MdTf_II and 6.5% of L1MdTf_III. In addition, 1,001 L1MdTf shorter elements were predicted as targets, as well as 22.5% of L1MdGf_II FL elements (69 elements) (Fig.S1B). The majority (74.5%) of L1MdTf elements were predicted to contain more than one binding site for the LKF repressor (Fig.S1B). Off-target sequences outside L1 repeats were found only at 6 genomic locations, including 1 located 3 bp away from an annotated L1MdTf element (Table S1). LKF and AKF repressor sequences were cloned upstream of an IRES-ZsGreen reporter, enabling a readout of expressing cells, under the control of doxycycline (DOX)-responsive promoter and knocked-in at the *Rosa26* locus in a PGK12.1 mouse ESC line (PGK-G10) (Fig.1A). Several independent clones were analysed by RT-qPCR and showed robust induction of the engineered repressor transgene (Fig.S1C). In addition, using primers specific for the 5’UTR region, we observed a 2-fold down-regulation of L1MdTf expression following 48h of DOX treatment in LKF-expressing clones only, and no changes in expression of L1MdA (Fig.1B).

**Figure 1.**
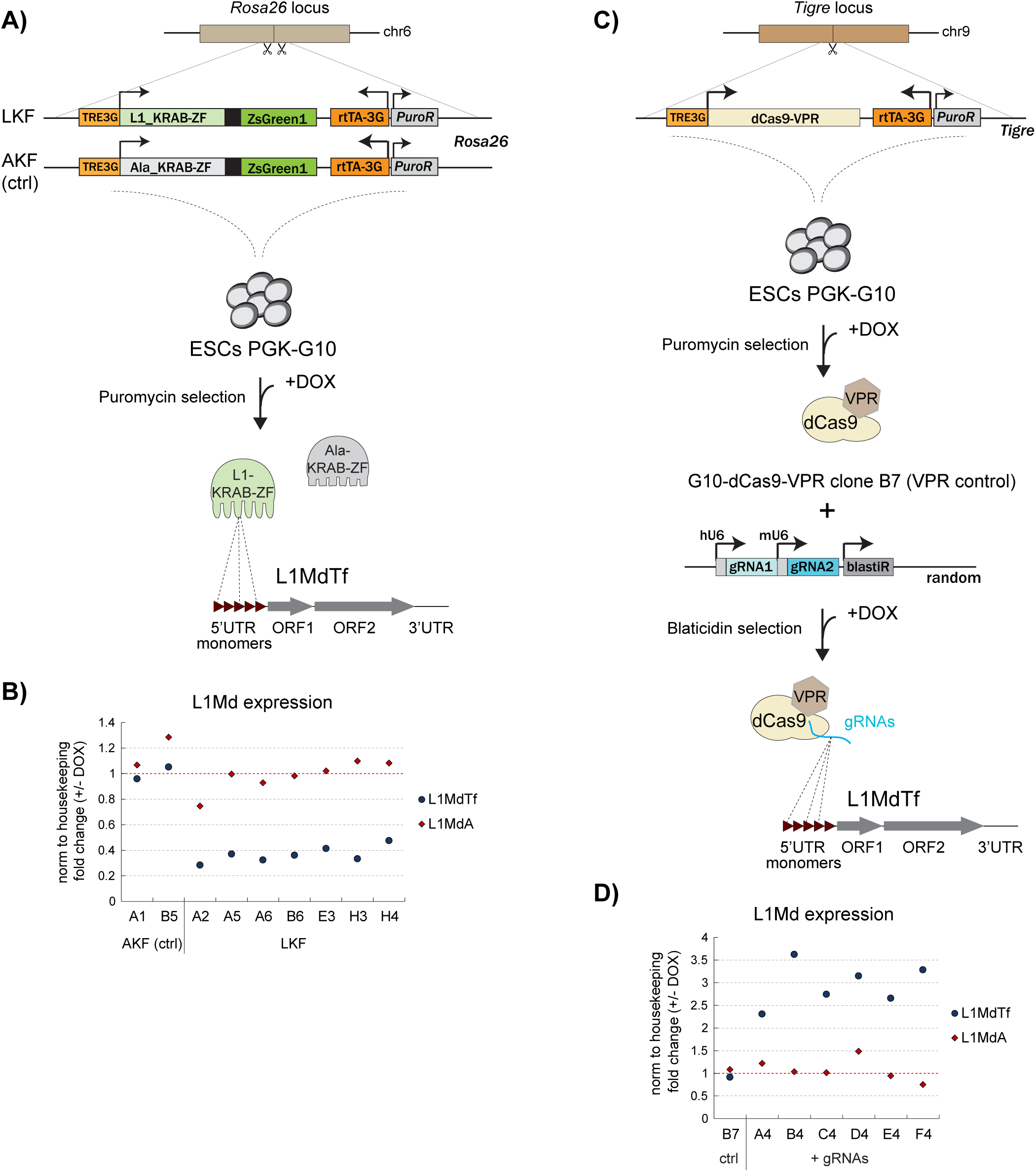
Targeting strategy for L1 expression perturbation in mouse ES cells. A) Structure of the DOX-inducible LKF/AKF transgenes transfected in PGK-G10 cells for knock-in at *Rosa26*. B) Following puromycin selection, colony picking and genotyping, 9 clones were analysed by qPCR for expression of the L1MdTf/L1MdA families using primers specific for the 5’UTR region, upon 48h of DOX treatment. C) Structure of the DOX-inducible dCas9-VPR transgene transfected in PGK-G10 cells for knock-in at *Tigre*. D) Following puromycin and blasticidin selections, colony picking and genotyping, 7 clones, including the control VPR-B7, were analysed for expression of L1MdTf/L1MdA upon DOX treatment (48h) by qPCR. Scissors indicate the approximate positions of gRNAs. TRE3G: promoter Tet-responsive element 3rd generation. rtTA 3G: DOX-inducible transactivator 3rd generation. PuroR: puromycin resistance gene. BlastiR: blasticidin resistance gene. hU6: human U6 promoter. mU6: mouse U6 promoter. ZF: zinc fingers.

To achieve genome-wide activation of L1MdTf elements, we relied on dead Cas9 (dCas9) fused to the VPR activation domain system, associated with two sgRNAs targeting two 20 bp sequences from the L1MdTf monomeric consensus sequence (Fig.1C,D, Fig.S1A). Similarly to the repression system, we checked the prevalence of the gRNAs targets *in silico* and found a total of 1,060 FL-L1MdTf elements and 991 L1MdTf shorter copies predicted to be targeted by one or both gRNAs, accounting for more than 95% of L1MdTf_I and L1MdTf_II FL elements, the majority (93.9%) of which contain more than one gRNA-binding site (Fig.S1D). Off-target sites outside L1s were found at 17 locations, again all within TEs (Table S2). The dCas9-VPR transgene placed under the control of a DOX-inducible promoter was also integrated into the PGK-G10 ESC line. Robust induction of the transgene upon 48h of DOX treatment was assessed by RT-qPCR in independent clones, of which one was selected (VPR-B7, used as VPR-control in all subsequent analyses) for further introduction of the dual sgRNA-encoding transgene (Fig.S1C). Analysis of independent clones showed significant activation of 2.5 to 3-fold specific for L1MdTf elements upon DOX induction of the dCas9-VPR. L1MdA expression, on the other hand, was not affected (Fig.1D). These results indicate that the engineered repressor and activator targeting specific young L1 subfamilies lead to their general and robust repression or activation.

### Characterization of ESC lines encoding engineered L1MdTf-repressor and activator

Based on our initial RT-qPCR results (Fig.1), we selected two clones showing efficient repression (LKF-A5 & -A6) and activation (VPR-B4 & -D4) of L1MdTf elements for further characterization, as well as one control clone for each system (AKF-B5, expressing the control repressor and VPR-B7, expressing no sgRNAs). First, we tested the DOX treatment duration to achieve maximal expression of the AKF/LKF repressors by analysing the ZsGreen reporter using microscopy and flow cytometry (Fig.2A,B). The results showed that the expression was constant in the selected clones after 24h, 48h, and 96h of treatment. Therefore, for all subsequent analyses, we decided to consider a DOX treatment of 48h. We then analysed the repression of L1MdTf and L1MdA families by RT-qPCR. We confirmed that L1MdTf expression was significantly reduced in the LKF-expressing clones with DOX, whereas L1MdA expression was unaffected. Similar levels of repression were observed when using primers specific for the ORF2 region, able to recognize all L1 families (Fig.2C), indicating that more than half of transcriptionally active L1 elements in mouse ESCs appear represented by L1MdTf elements. We also examined the repression at the protein level by measuring ORF1 protein, encoded by intact full-length L1s, by Western blot and found that the levels of the top band corresponding to the variant p43 isoform are reduced in the LKF-expressing clones upon DOX treatment (Fig.2D). This isoform mainly corresponds to the ORF1 protein encoded by L1MdTf elements (Kolosha and Martin 1995), highlighting the specific repression of L1MdTf elements both at the RNA and protein levels and indicative of repression of non-intact and intact L1MdTfs.

**Figure 2.**
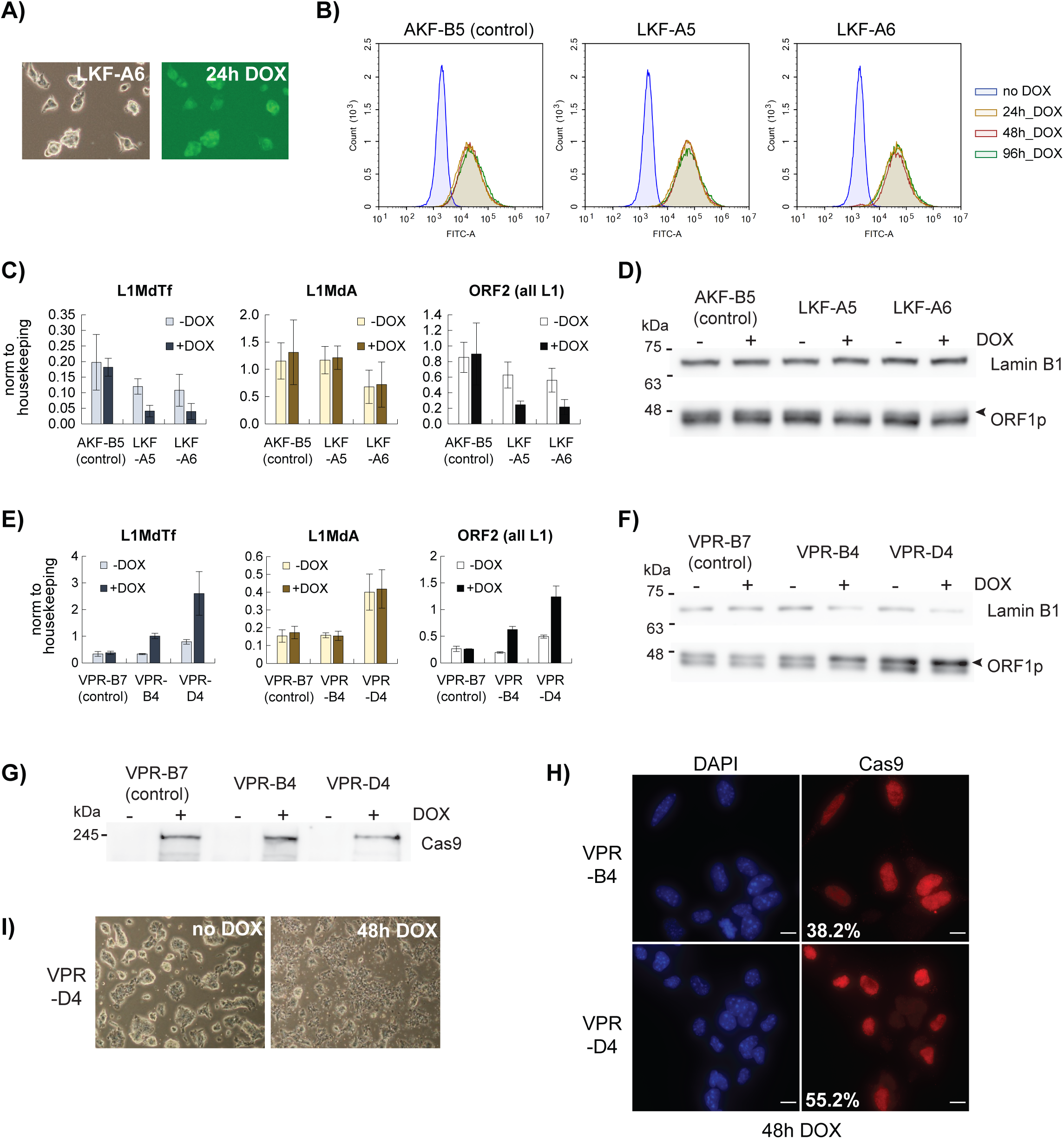
Characterization of ESCs showing L1MdTf expression perturbation. A) Brightfield and fluorescence microscopy images of ESC colonies from repressor clone LKF-A6 showing ZsGreen1 signal after 24h of DOX treatment. B) Flow cytometry analysis of repressor clones LKF-A5, LKF-A6 and control AKF-B5, showing similar fluorescence levels after DOX treatment for 24h, 48h and 96h. C) qPCR analysis of L1 expression using primers for the 5’UTR of L1MdTf/L1MdA and the ORF2 region of all L1s, showing specific repression of L1MdTf in repressor clones LKF-A5, LKF-A6 and not in control AKF-B5, upon DOX treatment for 48h. Levels are normalized relative to housekeeping gene expression. Error bars represent the calculated standard deviation considering 5 biological replicates. D) Western blot showing decreased ORF1p protein levels (top isoform, indicated with an arrow) in repressor clones LKF-A5 and LKF-A6, after DOX treatment for 48h. LaminB is used as loading control. This experiment was repeated twice with similar results. E) qPCR analysis of L1 expression using primers specific for the ORF2 region and the 5’UTR of L1MdTf/L1MdA, showing specific activation of L1MdTf in activator clones VPR-B4, VPR-D4 and not in control VPR-B7 upon DOX treatment for 48h. Levels are normalized relative to housekeeping gene expression. Error bars represent the calculated standard deviation considering 3 biological replicates. F) Western blot showing increased ORF1p protein levels (top isoform, indicated with an arrow) in activator clones VPR-B4 and VPR-D4, after DOX treatment for 48h. LaminB1 is used as loading control. This experiment was repeated twice with similar results. G) Western blot showing induction of Cas9 after 48h of DOX treatment in control VPR-B7 and activator clones VPR-B4 and VPR-D4. H) Representative images of immunofluorescence showing induction of Cas9 after 48h of DOX treatment in activator clones VPR-B4 and VPR-D4 (scale bars, 10 μm). Percentages indicate the number of cells with both high and low Cas9 signals. I) Brightfield images of ESC colonies from activator clone VPR-D4 showing increased cell death after DOX treatment for 48h.

We also performed RT-qPCR and Western blot analyses to confirm the specific activation of L1MdTf elements using dCas9-VPR. RT-qPCR of the L1MdTf and L1MdA subfamilies, as well as using primers overlapping the ORF2 region, showed increased specific expression of L1MdTf and ORF2 in the two VPR clones expressing sgRNAs, VPR-B4 and -D4, upon DOX treatment (Fig.2E). Using Western blot analysis, we observed a mirror image to the one for L1MdTf repression, with the variant p43 isoform levels being more elevated in the VPR-B4 and -D4 clones upon DOX induction (Fig.2F). As expected, after induction, dCas9 protein could be detected in all clones by Western blot and by immunofluorescence (IF) (Fig.2G,H). Surprisingly, IF analysis showed that dCas9 expression was very heterogeneous in the 3 VPR clones analysed, with cells expressing dCas9 at high (∼20% of cells) or low levels or not expressing the protein (Fig.2H). This heterogeneity prompted us to look at L1 nascent RNA and ORF1 expression at the single-cell level using RNA FISH and IF. RNA FISH was performed using a probe corresponding to the FL-L1spa element (Naas et al. 1998; Chow et al. 2010). L1spa RNA FISH signal appears as discrete dots within the nucleus and strong foci located mainly in the cytoplasm. The percentage of cells with these foci, their abundance, and size increased upon DOX induction in the VPR-B4 and -D4 clones (Fig.S2A). ORF1 protein was detected as bright foci in the cytoplasm of a fraction of cells prior to DOX treatment. Consistently, these foci were more abundant and observed in more cells after DOX induction (Fig.S2B). Quantification of both L1 RNA and ORF1 protein signal intensity indicated a higher value in VPR-B4 and -D4 clones upon DOX treatment, indicative of the augmented transcriptional activity of L1 elements (Fig.S2A,B). Interestingly, dCas9-VPR-B4 and D4 clones showed increased cell death compared to the control VPR clone and the LKF repressor-expressing clones after 24h of DOX treatment, and this difference became more evident after 48h (Fig.2I). However, we did not notice any changes in ESC self-renewal, as previously reported upon L1-RNA degradation (Percharde et al. 2018).

### Engineered effectors affect expression of a large number of L1MdTf elements

To analyse genome-wide L1MdTf repression or activation and its impact, we performed 100 bp paired-end total RNA-seq (stranded, with rRNA depletion) on five of the previously characterised ESC clones: one control (AKF-B5) and one repressor clone (LKF-A6), plus one control (VPR-B7) and two activator clones (VPR-B4 & VPR-D4). For all cell lines, two biological replicates were sequenced, before and after DOX induction (Fig.3A). RNA-seq analysis was performed following a standard pipeline, using two mapping strategies to report only uniquely mapped reads or multi-mapped reads randomly assigned to one genomic location. The first strategy was used to examine transcriptional changes of genes and specific repeat elements, while the second strategy was used as a proxy to analyse the expression of different repeat families from the Repeat Masker database. Approximately 12.5% of reads in each library were mapped to multiple locations, of which an average of 20% corresponds to non-exonic repetitive sequences. No global significant changes were observed for the L1 family in the repression samples (Fig.S3A). However, consistent changes after DOX treatment were observed for L1s in the VPR activation system, as normalized counts for L1 sequences were significantly increased for the VPR-B4 and -D4 libraries with DOX (Fig.S3B). LTR/ERV and SINE/B2 families showed fluctuating counts, but this was uncorrelated with the repressor/activator expression or DOX treatment (Fig.S3A,B, Table S3). We then analysed the fold change of L1 repeats counts by subfamily (Fig.S3C, Table S4). For both L1 repression and activation, the most affected subfamilies were the evolutionarily youngest ones: L1MdTf_I and L1MdTf_II. The L1MdTf_III, L1MdGf_II and L1MdA_III subfamilies were also impacted but to a lesser extent (Fig.S3C). Importantly, this was consistent with our initial *in silico* prediction (Fig.S1B,C). We also analysed the fold-change over individual elements from each L1MdTf subfamily, for which uniquely mapped reads could be identified with confidence. In line with our prior qPCR analysis (Fig.2C,E), repression of L1MdTf_I and L1MdTf_II elements is about four-fold in the LKF-A6 clone, and their activation is around three-fold in the VPR-B4 and -D4 libraries relative to AKF-B5 and VPR-B7 controls, respectively (Fig.3B). L1MdTf_III elements were also affected, although to a lesser extent (Fig.3B).

**Figure 3.**
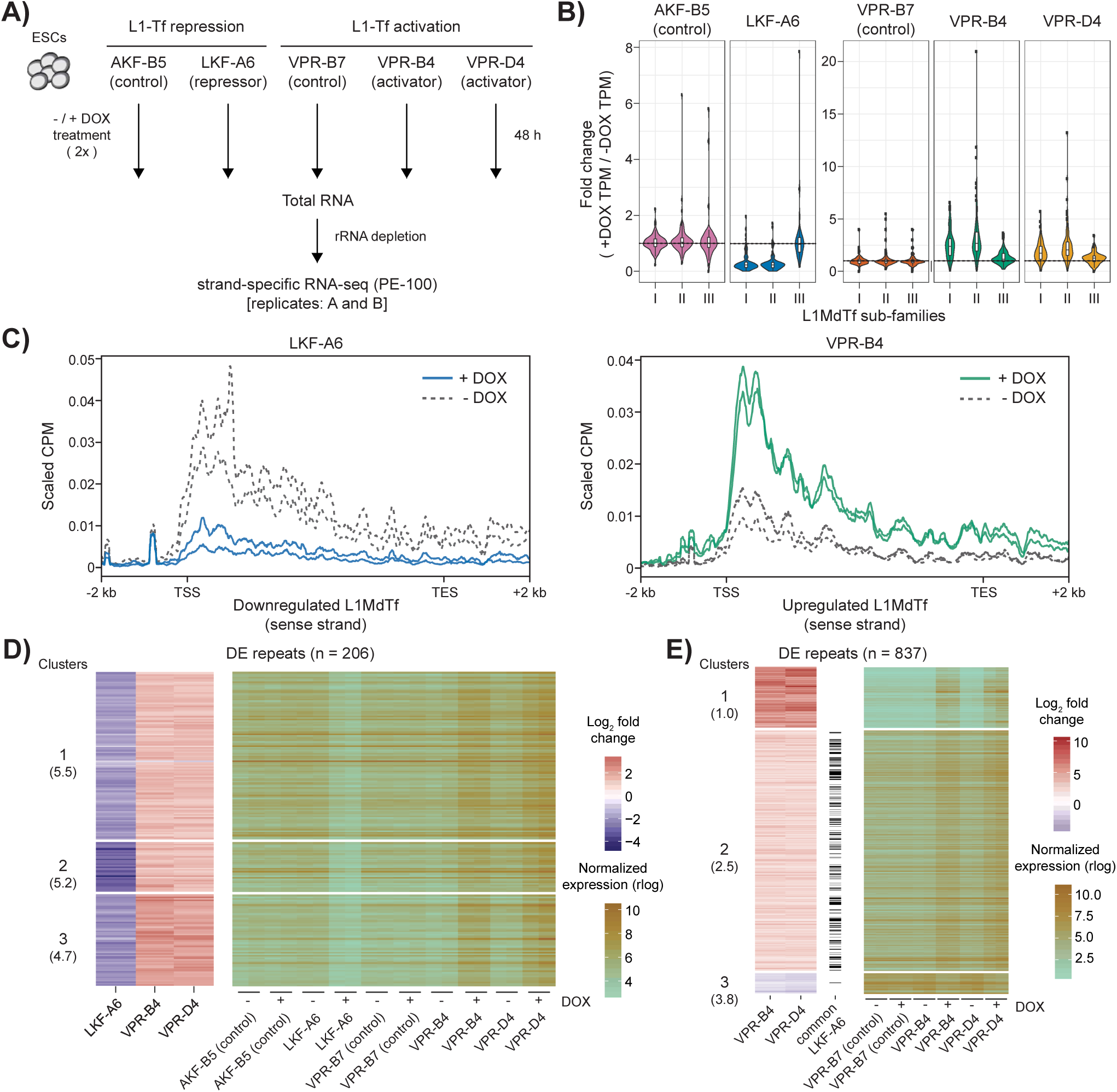
Differential expression of L1MdTf and other repeats upon L1MdTf repression and activation. A) Scheme of RNA-seq data generation from ESC selected clones treated with or without DOX. B) Violin plots representing the distribution of expression fold changes for individual L1MdTf repeats classified by subfamily in all clones. Dashed line indicates values equal to 1. C) Metagene plot of normalized RNA-seq signal (shown for LKF-A6 and VPR-B4, replicates A & B) on L1MdTf repeats showing downregulation in the LKF-A6 clone (left) or upregulation in the activator clones (right) upon DOX treatment. The x-axis is scaled to represent full annotations and the 2 kb upstream and downstream regions. D) Heat maps showing expression fold change (left) and normalized expression (right) of common DERs identified in repressor and activator clones. DERs are organized according to their cluster classification. Average normalized expression of genes in the -DOX condition of all displayed clones is indicated below each cluster number. E) Heat maps showing expression fold change (left) and normalized expression (right) of DERs identified in both activator clones, as described in D. Black bars indicate if the repeat is also identified as a DER in the LKF-A6 clone. CPM: counts per million reads mapped; TSS: transcription start site; TES: transcription end site.

To identify changes in the RNA-seq signal distribution along L1MdTf elements genome-wide, we determined their average sense and antisense profiles in all libraries using uniquely mapped reads. As some of the L1 annotations may not be perfectly delimited, we analyzed their expression using the annotated coordinates and a +2 kb window from the start and end of the element. Signal in sense direction was detected across the whole element, with higher coverage just downstream of the TSS, as expected because of the greater sequence diversity of the 5’UTR/promoter region among L1Md families. This trend was strongly enhanced by VPR induction and reduced upon expression of the repressor (Fig.3C; FigS4A). Interestingly, we also noticed read-through transcription after the 3’UTR towards downstream sequences (Fig.3C; FigS4A) and antisense transcription originating from the L1-promoter region towards upstream sequences in all samples (FigS4B). Both antisense and read-through transcription were affected by the engineered effectors upon DOX treatment, indicating that this is a direct consequence of L1 expression (Fig.3C; FigS4A,B), confirming prior observations regarding the existence of an antisense promoter in mouse L1 elements and a weak polyadenylation signal (Goodier et al. 2001; Li et al. 2014).

Differential expression analysis considering specific L1MdTf elements showed a significant correlation between the number of LKF or gRNA target sites in L1MdTfs and their expression fold change after DOX treatment, indicating that the simultaneous binding of dCas9 or the LKF repressor in the same region enhances activation or repression (Fig.S4C). Overall, using unique mapping, we found 513 individual repeat annotations differentially expressed (padj<0.01) in the LKF-expressing clone, with a large majority (n=377 out of 1,023 elements tested) corresponding to repressed L1MdTf annotations (Table S5). In the VPR-B4 and -D4 clones, we found 1,400 and 1,421 repeat annotations with significant changes in expression (padj<0.01), respectively, of which a large part corresponds to upregulated L1MdTf elements (n=699 out of 2,315 for VPR-B4, Table S6; n=656 out of 1,815 for VPR-D4, Table S7). Importantly, there is an extensive overlap (n=837) between differentially expressed repeats (DERs) in the two VPR clones (Fig.S4D), indicating that the effects observed are reproducible. By comparing the DERs in all clones, we identified 206 repeat annotations showing reciprocal expression changes mediated by the engineered effectors and corresponding mostly to L1MdTf elements (Fig.3D; Table S8). Clustering these repeats by their fold change showed that the L1MdTf repeats with stronger repression in the LKF-A6 clone (cluster 2) do not correspond to the L1MdTf repeats with stronger activation in the VPR clones (cluster 3), possibly due to the different effector systems that were used or the initial expression levels in ESCs. Indeed, the more strongly activated repeats (cluster 3) tend to have lower normalized expression prior to DOX treatment (Fig.3D). We also observed this trend when clustering the common DERs in the VPR clones. However, in this case, the cluster included both L1MdTf repeats and members of other L1 subfamilies (cluster 1) (Fig.3E; Table S9). As only a few gRNA target sites do not overlap L1MdTfs, this result suggests that increased expression of non-L1MdTf L1 repeats may be a secondary effect of L1MdTf upregulation rather than direct activation mediated by the dCas9 effector. The largest cluster of DERs in the VPR clones includes mainly L1MdTf_I and L1MdTf_II annotations that show higher basal expression levels and are repressed in the LKF-A6 clone (cluster 2). Importantly, we also identified a small fraction of downregulated repeats characterized by their high basal expression and corresponding to ERVK/ERVL annotations (cluster 3) (Fig.3E; Table S9). Altogether, these results indicate that the effector systems allow the specific modulation of L1MdTf expression genome-wide and suggest L1MdTf roles in the regulation of other repeat subfamilies.

### L1MdTf misregulation influences nearby host gene expression

By displaying the RNA-seq signal profiles, we observed that expression changes of L1MdTf DERs coincide with changes in the expression of genes in their proximity, as exemplified by the *Iqub* and *Pkhd1* genes which both contain intronic L1MdTf elements (Fig.4A). Therefore, we evaluated the impact of the genome-wide perturbation of L1MdTf elements on host gene expression using our RNA-seq datasets. For the repression system, we observed a mild impact on gene expression, with 91 differentially expressed genes (DEGs) (adjusted p-value (padj)<0.01) between the DOX and no DOX conditions for the LKF-expressing clone, of which 10 are also DEGs in the control repressor clone, thus likely sensitive to the DOX treatment alone. Among the 81 remaining genes, the majority (n=65) are downregulated (Fig.4B; Fig.S5A; Table S5; Table S10). For the activation system, the impact on gene expression was much stronger, with 1,384 and 1,457 DEGs (padj<0.01) between DOX and no DOX conditions for the VPR-B4 and -D4 clones, respectively (Fig.4B). A minimal number of DEGs were also detected in the control activator clone (<=14) and were excluded from all further analyses (Fig.S5A; Table S11). Of note, 31 DEGs were common between the repression and activation systems, with almost half of the downregulated genes in the LKF-A6 clone (29/65) showing upregulation in the VPR clones, as illustrated by the *Iqub* and *Pkhd1* genes (Fig.4A,C; Table S12). In the VPR clones, similar to what we observed previously for the DERs, the majority of DEGs upon DOX treatment are found in both clones (n=1,024) and are upregulated (n=923) (Fig.4B; Fig.4D,E; Table S13). We found that, overall, affected genes do not show apparent clustering and are located across all chromosomes. There were, however, some notable exceptions, including the X chromosome that showed a two-fold enrichment for upregulated genes (105 DEGs) compared to autosomes (43 DEGs on average), correlating with the two-fold enrichment of L1 repeats on the X (number of L1MdTf >6 kb in chrX=162; average number of elements in other chromosomes=79). Additionally, we observed the upregulation of small clusters of family-related genes (e.g. GABA receptor subunit, Keratin, Mage) and a large cluster of homeobox genes located on the X chromosome, the *Rhox* cluster (Fig.S5B). Similarly to DERs, genes with higher fold change upon VPR expression displayed lower expression levels in the -DOX condition (cluster 1 & 2) (Fig.4E). We selected eight genes among the top differentially expressed, including DEGs showing reciprocal repression or activation as well as X-linked genes, and performed RT-qPCR analysis. We confirmed the observed expression changes in both directions upon DOX treatment, including independent biological replicates (Fig.S5C,D).

**Figure 4.**
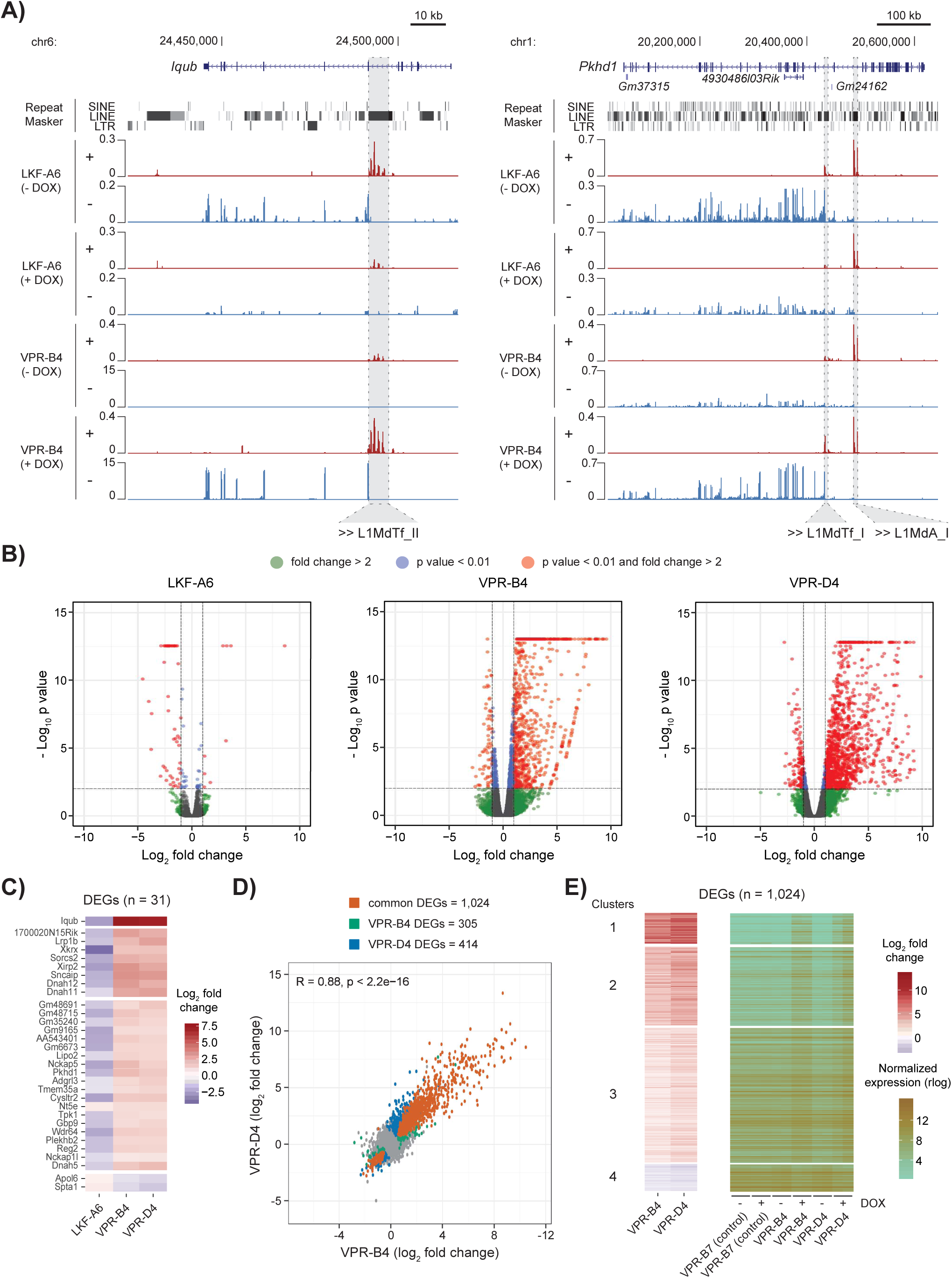
Differential expression of genes upon L1MdTf repression and activation. A) Examples of downregulated and upregulated genes following the repression and activation of L1MdTf repeats, respectively. RNA-seq profiles by strand are shown in red (+ strand) and blue (- strand) for the DOX-treated and untreated samples. Intronic L1MdTf repeats with reciprocal expression changes and a L1MdA repeat with no expression changes, and their orientation, are highlighted to exemplify the specific targeting. B) Volcano plots showing gene expression fold changes (x-axis) and their significance (y-axis) in the repressor and activator clones. C) Heat map depicting the fold changes of common DEGs identified in the repressor and activator clones. Genes are clustered based on their fold changes. D) Comparison of expression fold changes of genes in the VPR-B4 and VPR-D4 clones after DOX treatment. Grey dots represent genes with no significant changes in expression and colored dots represent genes identified as DEGs in both clones or in one of the clones only. Pearson correlation coefficient and significance are indicated. E) Heat maps showing expression fold change (left) and normalized expression (right) of common DEGs identified in both activator clones. DEGs are organized according to their cluster classification.

Given our observation that activation and repression of L1MdTf elements appear correlated with activation/repression of nearby genes (Fig.4A), we next examined the genomic distance between DEGs or DERs and misregulated L1MdTf elements. We first plotted the distance distribution between DE L1MdTfs and DEGs or DERs. Overall, we found that the average distance values of LKF-downregulated (∼3.9 Mb) and VPR-upregulated (∼3.9 Mb) genes to DE L1MdTf elements are lower than those of LKF-upregulated (∼5.6 Mb), VPR-downregulated (∼6.3 Mb), or genes whose expression remains unaffected (Fig.5A,B; Table S14; Table S15; Table S16). Moreover, this tendency was clearer for commonly upregulated genes in both VPR clones (∼3.1 Mb) than for genes upregulated only in one of the 2 clones (∼4.2 Mb) (Fig.5B). Similar results were observed for non-L1 DERs down- or upregulated in LKF and VPR clones, respectively (Fig.S6A,B; Table S17). Additionally, when comparing the fold change and the genomic distance between affected L1s and misregulated genes (for both LKF and VPR clones) or non-L1 DERs (for the VPR clones only), we observed that the genes or repeats showing the highest fold change upon DOX (the 90th percentile) tend to be located within an average distance <3 Mb from a L1MdTf DER (Fig.5C; Fig.S6C). This rather large average distance may be due to the fact that we can only analyse a limited number of specific L1MdTf using uniquely mapped reads due to the repetitive nature of these sequences, and probably, the proportion of DEGs and non-L1 DERs located close to affected L1MdTf repeats is underestimated. In fact, when comparing the fold changes of DEGs and the distance to all annotated L1MdTf elements, we observed that most of the strongly LKF-downregulated and VPR-upregulated genes are located within an average 260 kb distance from an L1MdTf element (Fig.S6D). For this reason, we tested if, at shorter distances (<=200 kb), the presence of a DE L1MdTf correlates with expression changes of genes and non-L1 repeats. This analysis showed significant differences from expected in the distribution of genes and repeats classified according to their expression changes and distance to DE L1MdTf elements (p-value < 1×10^-3, chi-square test of independence) (Fig.S6E,F). Indeed, most of these differences were driven by the downregulated (30.9% and 30.3% (values relative to DE only)) and the commonly upregulated (35.9% and 9.2%) genes and repeats in the LKF and VPR clones, respectively. In summary, our analysis shows that misregulation of L1MdTf elements tends to impact nearby transcription and gene expression. Further exploration of the genome tracks for specific genes adjacent to misregulated L1s suggest that these elements may act as short-range *cis*-regulatory elements (e.g., *Magee2*, *Fabp7/Smpdl3a* and *Ret* genes (Fig.S7A-C)) or as alternative promoters when located within intronic sequences (e.g., *Iqub*, *Pkhd1* and *Slc8a1* genes (Fig.4A; Fig.S7D)).

**Figure 5.**
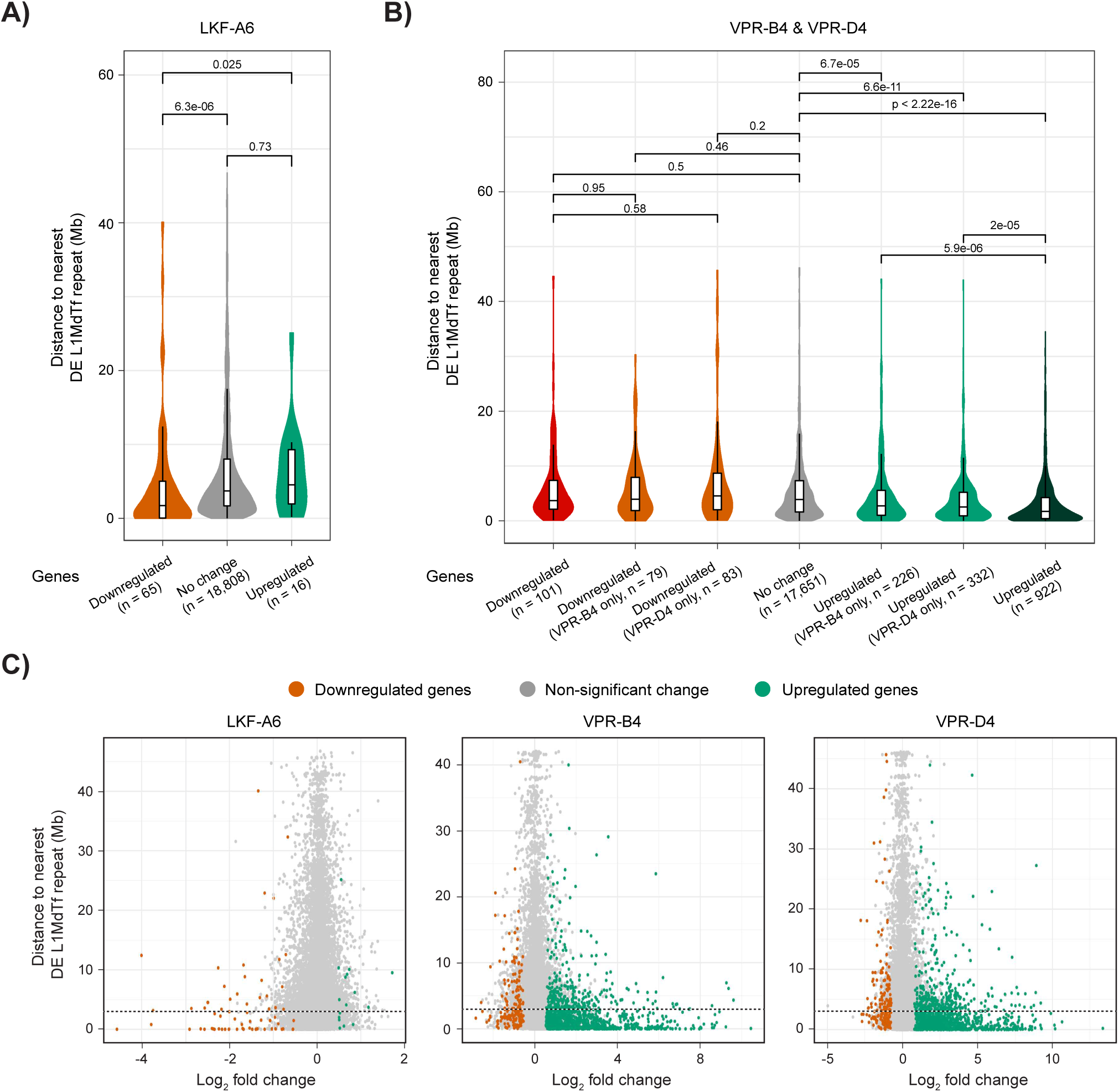
Distance analysis between DEGs and DE L1MdTf repeats. A) Distribution of distances between DE L1MdTf repeats and genes classified according with DE analysis results in the LKF-A6 clone. Significance values of the differences between distributions are displayed above violin plots and were calculated using Wilcoxon rank-sum tests. B) Distribution of distances between DE L1MdTf repeats and genes classified according with DE analysis results in the activator clones. Significance values as described for panel A. C) Comparison between expression fold change upon L1MdTf repression (LKF-A6) or activation (VPR-B4, VPR-D4) and distance to nearest DE L1MdTf repeat for all genes tested for DE. Dashed lines indicate a distance of 3 Mb.

Moreover, considering that the L1MdTf promoter region can drive transcription in both sense and antisense direction (Fig.3C; S4A,B), L1MdTfs could affect expression of genes, regardless of their orientation. For instance, we identified a transcript that starts within the L1MdTf_II element located upstream of *Iqub* exon 7 by RT-PCR (Fig.S7E), leading to the production of a chimeric transcript and indicating that the antisense promoter activity of the L1 element can drive expression of an *Iqub* transcript from exon 7. Interestingly, L1-mediated gene expression of *Iqub* could also occur in cells in the absence of DOX since the proximal exons of the gene appear expressed at lower levels than exons located after the L1 element (Fig.4A; Fig.S7E). As the L1MdTf promoter drives transcription in both directions (Fig.3C; S4B), we also investigated the orientation of the intronic DE L1MdTf elements with respect to the orientation of genes in which they are located, to determine the most prominent scenario. Out of the 53 upregulated and intronic L1MdTfs identified in the VPR clones, 31 were in the antisense orientation with respect to the upregulated genes. Similarly, 9 out of the 14 downregulated L1MdTfs overlapping downregulated genes in the LKF clone were also in antisense orientation. These proportions are similar to previous estimates of the orientation bias for intronic L1s (Nellaker et al. 2012). There is thus a higher proportion of antisense L1MdTf located within intronic sequences of genes (e.g., *Iqub* and *Pkhd1* in Fig.4A; *Slc8a1* in Fig.S7D), as expected given that repeats in the opposite orientation are less detrimental than sense-oriented ones (Zhang et al. 2011).

### Identification of elements and genes with skewed allelic expression

Taking advantage of the fact that the PGK-G10 cell line used to generate the L1MdTf repressor and activator lines is a hybrid F1 line with PGK and 129 genotypes (Penny et al. 1996; Marks et al. 2009), we sought to determine if the transcriptional changes observed for L1MdTfs occur specifically for one of the alleles. This could suggest that the associated repeats are polymorphic and present on only one allele. To assess this, we first identified repeats with skewed allelic expression. Considering all repeat families, more than 35 elements showed skewed expression towards each of the alleles in the two effector cell lines (Fig.S8A). These numbers are lower for the activator as only the repeats with significant differences from biallelic expression common between VPR-B4 and VPR-D4 are considered. No L1MdTfs were found among the repeats with skewed allelic expression, which is consistent with the fact that young L1s have low levels of sequence diversity (Teissandier et al. 2019). Actually, very few L1MdTfs (<10 elements in each of the comparisons) have uniquely mapped reads that could be assigned to one of the alleles. Nevertheless, as an alternative strategy to identify L1MdTfs affecting gene expression in *cis* on one allele only, we next identified genes with skewed expression and checked if they corresponded to DEGs upon perturbation of L1MdTfs. Overall, more genes (268 for the repressor and 210 for the activator) than repeats (135 for the repressor and 82 for the activator) showing skewed allelic expression were identified for the two alleles (Fig.S8B). However, only 14 and 1 were DEGs in the L1MdTf activator and repressor clone analyses, respectively. Two examples of such genes are *4930579G24Rik* and *Cysltr2*, which are located proximally to L1MdTfs, 1,336 bp and 88,908 bp away, and show skewed expression for the PGK or 129 allele respectively (Fig.S8C).

In this analysis, we assigned RNA-seq reads to alleles using only those SNPs that can be annotated with high confidence as PGK or 129 based on public annotations (see methods). Therefore, future work to complement the assignment of all heterozygous SNPs detected or using hybrid lines with different genetic background will improve the detection of L1MdTfs with allele-specific functions.

### Misregulated genes by L1MdTf effectors are enriched at repressed regions and are part of the neuronal network

To appreciate further the role that young L1s may play in gene regulation, we investigated whether DERs and DEGs found in the LKF and VPR clones (Table S5-S7) are located in specific genomic or chromatin environments using published ESC ChromHMM state annotations. These annotations classify mouse genomic regions in 12 states based on 7 chromatin marks (H3K27me3, H3K36me3, H3K27ac, H3K4me1, H3K4me3, H3K9ac, H3K9me3) and binding profiles of 3 transcription factors (CTCF, Nanog, Oct4) (Pintacuda et al. 2017). First, we looked at L1s and found that VPR-upregulated and LKF-repressed L1 elements tend to be associated with chromatin features of active promoter and intergenic regions (Fig.6A). Enrichment at active promoters was consistent with our observation that misregulated L1s could act as promoters, controlling nearby gene expression. Of note, VPR-upregulated L1s showed general enrichment in such regions independently of their expression fold change, as reflected by the common results obtained for the repeat clusters 1 and 2 (Fig.3E; Fig.S9A). We next analysed the chromatin features associated with non-L1 DERs. We found that VPR-upregulated and LKF-downregulated non-L1 repeats colocalize mainly with heterochromatin, but also with promoter or poised enhancer regions (Fig.6A). VPR-downregulated repeats on the other hand, corresponding mainly to repeat cluster 3 and ERVs elements (Fig.3E), are enriched with chromatin marks found at strong enhancers (FigS9A). In the case of DEGs, we observed that weakly and strongly VPR-upregulated DEGs are enriched in bivalent promoter chromatin domains, as well as repressed chromatin and intergenic regions (Fig.6A; Fig.S9A). Genes associated with bivalent promoters or located in repressed/intergenic chromatin environments have generally low expression levels or are silenced in ESCs (Bernstein et al. 2006). This is therefore in agreement with our observations that a fraction of VPR-upregulated genes, found in cluster 2 and 3 DEGs, tend to be weakly expressed in ESCs (Fig.4E). Some of these genes may be silenced or poised for subsequent activation later during ESC differentiation. On the other hand, VPR-downregulated genes (cluster 4 (Fig.4E)) colocalize with enhancer chromatin states, as observed for downregulated non-L1 repeats (Fig.6A; Fig.S9A). Finally, we mainly observed LKF downregulated genes at intergenic regions, possibly due to the lower number of regions considered (n=65). Overall, our observations highlight that most genes misregulated following L1 perturbation are found within repressive or silenced domains in ESCs.

**Figure 6.**
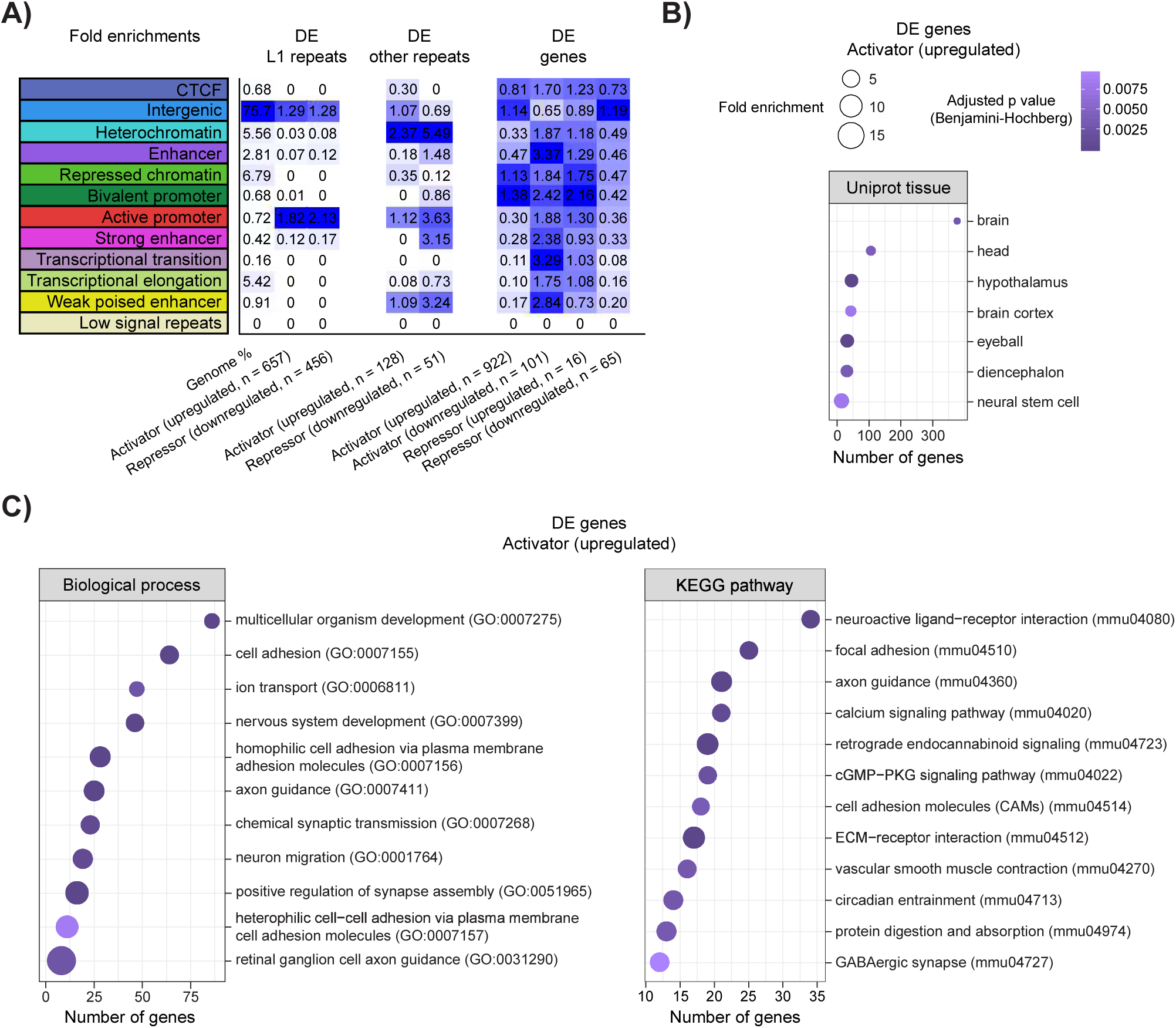
Chromatin state and GO enrichment analyses for DEGs and DERs. A) Heat map showing fold enrichment for each chromatin state of DE repeats and genes classified based on expression fold change direction. Increased color darkness represents increased enrichment. B) Expression enrichment of common upregulated genes in activator clones relative to Uniprot tissue terms. The number of genes by term is shown on the x-axis, circle size indicates fold enrichment and darker colors indicate higher significance. C) GO enrichment analysis of biological process terms (left) and KEGG pathways (right) for the common upregulated genes in activator clones. Plot annotation as described for panel B.

We next performed a gene ontology and KEGG pathway enrichment analysis to identify specific terms or pathways enriched in our differentially expressed gene sets. Given that the number of DEGs was too low in the repression system (n=81), this analysis was only performed for upregulated (n=923) and downregulated (n=101) genes in the activation system (Fig.6B,C; Fig.S9B-D).

Interestingly, we found that metabolism related terms are enriched in the downregulated gene set (Fig.S9B; Table S18). This could be related to the increased cell death observed following DOX induction and L1MdTf activation. For the set of upregulated genes, we first observed enrichment for genes specifically expressed in brain regions, tissues, and neural stem cells (Fig.6B). Biological process and KEGG pathway analysis revealed a predominance of neuronal function and development-related terms (Fig.6C; Table S19). Interestingly, analysis of cluster 2 and cluster 3 DEGs (Fig.4E; Table S20; Table S21) revealed an over-representation of genes expressed at the cell surface and related to synaptic function (Fig.S9C,D). Most of these genes are not expressed or have low expression levels in ESCs compared to other upregulated genes that are not part of these processes and pathways (Fig.S9E). As the activity of L1 elements is reportedly higher in neural stem cells and neurons, our data suggest that the expression of these sets of genes may be correlated with higher expression of L1.

## Discussion

In this study, we used engineered effectors to perturb the expression of full-length L1 elements and test their regulatory potential in mouse ESCs, focusing on one of the three most active L1Md subfamilies in the mouse genome, independently of their coding nature and retrotransposition competence. Our data show that perturbing FL-L1MdTf expression in ES cells has an impact on the expression of genes or repeats located in *cis* in the neighboring environment of these elements. Remarkably, L1MdTf activation is associated with a much higher number of misregulated genes than L1MdTf repression. Moreover, our analysis revealed that most of the affected genes, upon L1MdTf activation, are silenced or lowly expressed genes in ESCs, and a significant fraction of these have neuronal-related functions.

Previous analyses addressing the effects of interfering with mouse L1 transcriptional activity using TALE technology or ASOs for L1 RNA knockdown reported an impact on chromatin accessibility. However, in both studies, the authors found no evidence of misregulation of genes with an L1 element located within or in close proximity (Jachowicz et al. 2017; Percharde et al. 2018). The differences with our study may result from the technology used to interfere with L1 expression, the identity of targeted L1 elements, or the model used. TALE-mediated activation or repression of FL-L1 elements, from various subfamilies, was performed in mouse preimplantation embryos. The authors found that perturbing L1 transcriptional activity affects global chromatin condensation, with minor consequences on gene expression, including genes located near L1 elements (Jachowicz et al. 2017). ASOs targeting the coding region of L1s were used in mouse ESCs and preimplantation embryos to induce the depletion of L1 transcripts produced from any L1 subfamily (Percharde et al. 2018). Unlike CRISPR or TALE-based methods that interfere directly with transcription, this approach enables the analysis of exclusively *trans*-regulation effects mediated by L1 RNA or eventually L1-encoded proteins. The latter is exemplified by the finding that L1 RNA functions as a nuclear RNA scaffold in mESCs repressing essentially the 2-cell embryo transcription program. Nevertheless, expression of genes located proximally to L1s was not affected (Percharde et al. 2018).

The engineered effectors used in our study affect transcription and hence lead to a reduction of global L1 RNA and protein levels, as shown by qPCR, RNA FISH, and western blot analysis. This strategy allows us, therefore, to test both the *cis*- and *trans*-effects mediated by L1- encoded products. We did not observe a loss of ESC self-renewal nor activation of the 2-cell gene expression program following L1 repression, as previously reported following L1 RNA depletion (Percharde et al. 2018; Lu et al. 2020; Wei et al. 2022). The extent of total L1 RNA levels depletion might explain the differences observed, given that in this study we are targeting only one active L1 subfamily. We noticed, however, that one of the two VPR clones (VPR-D4) analysed showed significant upregulation of some genes belonging to the 2-cell program (e.g., the cluster of *Zscan4* genes on chr7). This was not observed in the other clone (VPR-B4) or to a much lesser extent, which may reflect cell to cell differences existing in the population prior to subcloning. It should be noted that these genes do not appear to lie near misregulated FL-L1. These observations suggest an indirect effect, whereby perturbation of L1 RNA levels, whether increased (our study) or decreased (Percharde et al. 2018; Lu et al. 2020; Wei et al. 2022), has an impact on chromatin organization with consequences on gene expression, as previously suggested (Percharde et al. 2018; Lu et al. 2020; Wei et al. 2022). Genes from the 2-cell stage expression program appear to be particularly sensitive to changes in chromatin structure. We hypothesize that other gene clusters found to be affected in both VPR clones, such as GABA receptor subunit, Keratin, Mage, *Rhox* genes, may also be particularly sensitive to changes in chromatin organization following perturbation of L1 RNA levels.

In our study, we found that a significant proportion of genes or repeats showing differential expression following L1MdTf expression perturbation, are located within a <3 Mb distance from misregulated FL-L1MdTf. This finding suggests that these elements may act as enhancers when located in intergenic regions (Fig.S7A-C) or alternative promoters when located in introns (Fig.4A; Fig.S7D,E). This was corroborated by our ChromHMM state analysis, showing that a proportion of misregulated L1MdTfs tends to colocalize with chromatin signatures of active promoter regions. Interestingly, a recent analysis in human ESCs showed that the 5’UTR regions of some active young L1 elements are enriched with H4K16ac and H3K122ac, chromatin features associated with active enhancers (Taylor et al. 2013; Pradeepa et al. 2016; Pal et al. 2023). Downregulation or deletion of acetylation positive- L1s, but not of acetylation negative-L1s, is associated with downregulation of genes in *cis*. This indicates that these elements, which can be found within introns or in intergenic regions, can function as *cis*-regulatory elements, a scenario that echoes our observations. It remains to be seen whether specific chromatin signatures could be defined for those active mouse L1s elements that were found to influence nearby gene expression in mESCs. Both our study and these recent findings collectively reveal that a subset of young and active L1s elements, in both mouse and human cells, have *cis*-regulatory functions.

Importantly, we observed that both sense or antisense transcription activity of L1MdTfs could modify nearby gene transcription or lead to the production of chimeric transcripts. This was previously reported for L1 elements in human cells, where the antisense promoter (ASP) of full-length L1s has been well characterized (Speek 2001). The ASP was shown to act as an alternative promoter for more than 40 protein-coding genes (Speek 2001; Matlik et al. 2006). Interestingly, both the canonical sense and antisense L1 promoters appear expressed in a tissue-restricted fashion in human tissues, suggesting that L1 activity might broaden the transcriptional potential for a given gene (Matlik et al. 2006; Faulkner et al. 2009). Our study uncovers that this could extend to mouse cells, especially considering the transcriptional dynamics of the mouse L1 promoter during early development, in neuronal lineages and certain disease contexts. Furthermore, our allelic expression analysis illustrates how polymorphic TE insertions can contribute to strain-specific variation in gene expression, as highlighted in a recent genomic analysis of 20 laboratory mouse strains (Ferraj et al. 2023). Our results suggest that some L1MdTfs insertions analysed in this study are potentially polymorphic in the ESC line used and can impact gene expression in an allele-specific manner. Of note, the reported number of genes showing skewed allelic expression is likely underestimated due to the limited number of high confidence SNPs used.

In terms of previously identified putative targets of TE-mediated gene regulation, the case of the *Fabp7* gene is a noteworthy example*. FABP7* ectopic expression was shown to be controlled by an LTR2 promoter located 30 kb upstream in a subset of human diffuse large B cell lymphomas, resulting in the expression of a TE-gene chimeric transcript (Lock et al. 2014). In our study, we observed that *Fabp7* and the neighbouring gene *Smpdl3a*, which are not expressed in ESCs, are upregulated following VPR induction. This is correlated with the upregulation of an L1MdTf_I element located 10 kb upstream and transcribed in the same direction (Fig.S7B). We speculate that the L1MdTf element may act as an enhancer controlling the expression of both genes located downstream rather than the production of chimeric transcripts. This could be yet another example where different species have independently co-opted specific TEs with regulatory potential to regulate the same gene (Chuong et al. 2017).

Our findings suggest that the transcriptional potential of young L1MdTf elements can be repurposed by the host genome in specific contexts or specific regions to regulate nearby genes in *cis*. This could be particularly relevant in developmental or tissue-specific contexts, where increased L1 activity has been observed. For instance, our GO analysis revealed that a significant proportion of VPR-activated genes have brain or neuronal-related functions. These genes are predominantly silenced or have low expression levels in ESCs, only becoming active later during differentiation. Considering that L1 elements activity was reported to be higher in neuronal cells, there could be a potential correlation between the coordinated expression of L1s and VPR-activated genes in neuronal cells. This raises the possibility that certain L1 elements have evolved to control gene networks within neuronal contexts. In addition, L1 transcription could particularly influence genomic regions where young L1 elements are more abundant, such as the X chromosome. Importantly, considering that these elements can become reactivated in certain diseases, such as cancer or neurodegenerative disorders (Burns 2017; Ravel-Godreuil et al. 2021), their misregulation could represent a significant mechanism through which TEs influence disease states. In the future, it will be important to use similar engineered effectors approaches to address whether and how TE expression could contribute to the establishment or progression of disease.

## Methods

### Genomic engineering of ESCs

#### ES cell lines

All cell lines used in this study are derived from feeder-independent PGK12.1 mouse female embryonic stem cells (ESCs) (Norris et al. 1994; Penny et al. 1996) and were maintained on gelatin-coated flasks in serum-containing ES cell medium (DMEM supplemented with 15% ESC-grade foetal bovine serum (FBS), 1000U/ml leukemia inhibitory factor (LIF, Millipore) and 0.1mM B-mercaptoethanol (Sigma)) in a humidified atmosphere at 37°C with 8% CO2. PGK12.1 is a hybrid cell line, which originates from mouse crossbreeds of two inbred strains (129 and C3H/He; both Mus musculus domesticus) with the distant strain PGK-1a/Ws (Mus musculus musculus) (Penny et al. 1996; Marks et al. 2009).

The PGK-G10 line employed in this study for genome engineering (derived from PGK12.1) exhibits a heterozygous deletion spanning a 5 kb segment encompassing the Xist promoter on chromosome X, ranging from the Xist TSS to 5 kb upstream (unpublished).

#### Plasmids

Knock-ins of the PGK-G10 ESC line were generated via CRISPR/Cas9 mediated homologous recombination. Donor plasmids used for knock-ins at the *Rosa26* (pEN111) and *Tigre* locus (pEN366, Addgene #156432), as well as the *Rosa26*- and *Tigre*-specific sgRNA-encoding plasmids (pX335-EN479, pX335-EN481 and pX330-EN1201 (Addgene #92144)) were all provided by E. Nora (UCSF).

The L1MdTf binding-zinc fingers were designed to bind a 20 bp sequence found within the consensus monomer sequence from the 5’UTR region of L1MdTf elements (Fig.S1A). A DNA fragment containing an array of 6 L1MdTf zinc-fingers, a KRAB repressor, and a FLAG tag sequences (LKF), as well as a DNA fragment comprising the mutated version (AKF), were then synthesised by GenScript and initially cloned into pLVX-IRES-ZsGreen1 lentiviral vector (Takara Bio) in frame with an IRES-ZsGreen1 sequence. The LKF/AKF-IRES-ZsGReen1 fragments (2138 bp) were amplified by PCR and cloned at the ClaI site in the *Rosa26*-targeting vector pEN111 by Gibson assembly (New England Biolabs). The resulting vectors obtained (pEN111-TRE3G-LKF/AKF-IRES-ZsGreen1-rtta3G_Rosa26-donor) were amplified and sequenced to ensure correct sequence and cloning.

The TRE3G-dCas9-VPR fragment was obtained by digestion of the PB-TRE-dCas9-VPR vector (Addgene #63800) with PmeI and PspXI. The fragment was subsequently cloned by DNA ligation in the donor vector pEN366 (Addgene #156432), after removal of the TRE3G-CTCF-mRuby2 fragment by digestion using the same restriction enzymes, PmeI and PspXI. The resulting vector obtained (pEN366-TRE3G-dCas9-VPR-rtta3G_Tigre-donor) was amplified and sequenced to ensure correct cloning.

Guide RNAs for L1MdTf elements were designed using the 212 bp consensus sequence of L1MdTf-monomers obtained from Goodier et al (Goodier et al. 2001) and the CRISPOR online tool (Concordet and Haeussler 2018). Two sgRNAs (sgTf-mono2 and sgTf-mono3) binding the monomer at two independent locations (Fig.S1A) and highly represented across L1MdTf elements in the genome were selected. The pLKO1-blast-U6-sgRNA-BfuA1-stuffer was used for the dual cloning of these two sgRNAs in tandem following the strategy described by (Holoch et al. 2021). The resulting construct, pLKO1-blast-U6_dual_sgTf-mono2-3, was amplified and sequenced to ensure correct cloning and sequence.

#### Transfections of ESCs and clone isolation

Transfections of DNA constructs in ESCs were performed using the P3 Primary Cell 4D-Nucleofector X Kit (V4XP-3024) and the Amaxa 4D Nucleofector system (Lonza). For each nucleofection, 5 million cells were resuspended in the nucleofection mix (prepared according to manufacturer’s instructions) and electroporated with 2.5 μg each of non-linearized vectors. Nucleofected cells were then serially diluted and plated on 10-cm dishes for colony picking and selection.

For knock-ins of the LKF or AKF transgenes at *Rosa26*, the PGK-G10 cell line was nucleofected with pEN111-TRE3G-LKF-IRES-ZsGreen1-rtta3G_Rosa26-donor (or AKF), pX335-EN479 and pX335-EN481. Puromycin selection (1 μg/ml) was started 2 days after transfection and kept for 8-10 days. Single colonies were then picked into 96-well plates. Genomic DNA was isolated directly in 96-well plates for PCR-based screening of knock-ins using primers overlapping the inserted construct and the surrounding *Rosa26* region. Twelve successfully knocked-in clones were selected, amplified and analysed for induction of expression of the LKF/AKF transgene and the impact on repression of L1MdTf elements upon doxycycline (DOX) (1 μg/ml) treatment for 48h.

For knock-ins of the dCas9-VPR transgene at *Tigre*, the PGK-G10 cell line was nucleofected with pEN366-TRE3G-dCas9-VPR-rtta3G_Tigre-donor and pX330-EN1201. Puromycin selection (1 μg/ml) was started 2 days after transfection and kept for 8-10 days. Single colonies were then picked into 96-well plates. Genomic DNA was isolated directly in 96-well plates for PCR-based screening of knock-ins using primers overlapping the inserted construct and the surrounding *Tigre* region. Four successfully knocked-in clones were selected, amplified and analysed for induction of expression of dCas9-VPR upon DOX treatment (1 μg/ml) for 48h. One dCas9-VPR-expressing clone (VPR-B7) was selected for transfection with pLKO1-blast-U6_dual_sgTf-mono2-3. Transfected cells were selected with blasticidin (10 μg/ml) 24h after transfection for 10-12 days. Cells were first analysed in bulk to test sgRNAs efficiency with or without DOX (1 μg/ml) by RT-qPCR. Cells were then seeded in 10-cm dishes to ensure optimal density for colony-picking in 96 well plates. Twelve clones were expanded and screened in 6-well plates in the presence or absence of DOX by RT-qPCR to assess L1MdTf upregulation efficiency.

Sequences of all sgRNAs and genotyping primers used for each engineered locus can be found in Table S22.

#### RNA extraction, reverse transcription, qPCR and PCR

For RNA extraction, cells grown in 6-well plates in the presence or absence of DOX (1 μg/ml) for 48h were briefly washed in 1X PBS and harvested directly in Trizol (Invitrogen). Total RNAs were extracted from Trizol with chloroform by phase separation. The aqueous phase was then mixed with an equal volume of 70% ethanol and transferred to a silica column from the RNeasy Mini kit (Qiagen). Total RNAs were then purified according to the instructions of the manufacturer, including on-column DNAse I digestion (Qiagen).

Total RNAs (500 ng to 1 µg) were reverse-transcribed into cDNAs using random primers and SuperScript III reverse transcriptase (Invitrogen) for 1 h at 50 °C. Quantitative PCR (qPCR) was performed using Power SybrGreen PCR master mix on a ViiA 7 real-time PCR system using standard settings (Applied Biosystems). Expression of genes was normalised to two housekeeping genes (*Bactin*, *Rrm2*). Standard PCR was performed on cDNAs using the GoTaq DNA polymerase (Promega). All primer sequences can be found in Table S23.

#### Western Blot analysis

Whole-cell extracts were prepared using RIPA buffer supplemented with protease inhibitors and proteins were separated by SDS-PAGE (8%) and blotted onto a nitrocellulose membrane using the dry transfer iBlot System (Thermo Fisher Scientific). Western blotting was performed using PBS-Tween 0.05%. Primary antibodies to ORF1p and LaminB1 were from Abcam (ab216324 (dilution 1:1000) and ab16048 (1:2000)). Primary antibody for Cas9 was from the Recombinant Antibodies Platform at Institut Curie (A-P-R#56, 1:2500). Secondary antibody anti-rabbit-HRP (Biorad, 170-6515) was used for ECL detection in accordance with the manufacturer’s recommendations using an Amersham imager 680.

#### Flow cytometry

Cells grown in 6-well plates in the presence or absence of DOX (1 μg/ml) for 48h were trypsinized, harvested in medium and then washed in 1X PBS before fluorescence analysis. ZsGReen fluorescence was analysed using the NovoCyte 2000R benchtop flow cytometer (Agilent). Fluorescence was acquired and analysed using the NovoExpress software following the manufacturer’s instructions. Briefly, percentage of ZsGreen-positive cells was determined upon definition of three gates: (i) FSC-H vs SSC-H to isolate cells from debris, (ii) FSC-H versus FSC-A to isolate single cells and (iii) FSC-H versus FITC-A for detection of ZsGreen-positive population.

#### Immunofluorescence

For IF, ES cells were grown on coverslips for 48h in the presence or absence of DOX (1 μg/ml). Cells were rinsed in 1X PBS, fixed for 10 min in 3.7% paraformaldehyde, rinsed in 1X PBS and permeabilised for 5 min in 1X PBS/0.5% Triton on ice. Cells were then blocked for 30 min at room temperature in 1X PBS/0.2% fish skin gelatin, and successively incubated with primary and secondary antibodies in 1X PBS/0.05% tween supplemented with 0.2% fish skin gelatin and 0.1% sodium azide for 1h at room temperature and washed after each incubation 3 times in 1X PBS/0.05% tween (5 min each). Cells were then counterstained with DAPI. Primary antibodies were as follows: rabbit anti-LINE-1 ORF1 (1:1000; Abcam, ab216324) and rabbit anti-Cas9 (1:100; Recombinant Antibodies Platform at Institut Curie, A-P-R#56). Secondary antibodies were goat anti-rabbit coupled with Cy3 (1:200; Bethyl, A120-201C3) and goat anti-mouse coupled with DyLight 488 (1:200; Bethyl, A90-116D2).

Images were acquired using Zeiss LSM 710 microscope system (ZEISS Microscopy) equipped with a Plan-Apochromat DIC 63x NA 1.40 oil objective and ZEISS ZEN software. The images were processed in FIJI using sum slices Z projection. For Cas9, the percentage of positive cells (with high or low signal) was manually counted in the population of cells treated with DOX. To quantify the L1-ORF1 signal, identical intensity threshold values were set in all the images of each clone or condition and the mean grey value was measured in 50-90 cells per clone/condition. The comparison of different conditions was based on the average of the mean grey values and their standard deviation. The statistical significance was assessed by two-tail t-Test.

#### RNA FISH

For RNA FISH, ES cells were grown on coverslips for 48h in the presence or absence of DOX (1 μg/ml). Cells were rinsed in 1X PBS, fixed for 10 min in 3.7% paraformaldehyde, rinsed in 1X PBS and permeabilised for 5 min in 1X PBS/0.5% Triton on ice, in the presence of 1% VRC (Vanadyl Ribonucleoside Complex; NEB, S1402S). Coverslips were then rinsed 3 times in 70% EtOH and stored at -20C. RNA FISH was performed essentially as previously described (Chaumeil et al. 2008). Briefly, coverslips, stored in 70% EtOH, were dehydrated in 80%, 95% and 100% EtOH (5 min each) and then allowed to air-dry. For L1 RNA detection, a full-length L1MdTf element (L1spa) cloned into pBluescript (TNC7; (Naas et al. 1998)) was used. Probes were labelled by nick translation (Vysis) with Spectrum GreendUTP following the manufacturer’s instructions and precipitated in the presence of salmon sperm. Labelled probes were denatured for 5 min at 74C. Cells were then directly hybridised with probes overnight at 37 C. After hybridization, the coverslips were washed 2 times in 50% formamide/2X SSC and 2 times in 2X SSC at 42C. Cells were counterstained with DAPI. Images were acquired using Zeiss LSM 710 microscope system (ZEISS Microscopy) equipped with a Plan-Apochromat DIC 63x NA 1.40 oil objective and ZEISS ZEN software. The images were processed in FIJI using sum slices Z projection. Identical intensity threshold values were set in all the images of each experiment and the mean grey value of TNC7 was measured in 80-200 cells per condition. The comparison of different conditions was based on the average of the mean grey values and their standard deviation. The statistical significance was assessed by two-tail t-Test.

#### In silico prediction of LKF and sgRNA target sites

Sites for the LKF repressor were searched in the mm10 mouse genome using bowtie (Langmead et al. 2009) (version 1.2.3) with options: -f -v 0 -y -a. To identify sgRNA sites, alignments to the sgTf-mono2 and sgTf-mono3 sgRNAs sequences without considering the PAM sequence were first identified using RIsearch2 (Alkan et al. 2017) (version 2.1) with the following options: -s 1:20 -m 2:0 -e 10000 -l 0 --noGUseed -p3. RIsearch2 results were used as input for CRISPRoff (Alkan et al. 2018) (version 1.1.2) to detect sites with an NGG PAM sequence with options: --evaluate_all --no_azimuth. LKF and sgRNA predicted sites were assigned to RepeatMasker annotations using bedtools intersect (Quinlan and Hall 2010) (version 2.29.0) with options: -wao -f 1. To quantify the number of target sites overlapping individual repeats, bedtools intersect was used with the option -c.

#### RNA-seq

All 5 selected ESC clones were sequenced in biological duplicates. For each sample, total RNA (800 ng) were used for library preparation using the Illumina TruSeq stranded total RNA Library Prep kit following the manufacturer’s protocols. Sequencing was performed in 100pb paired-end reads using a Novaseq 6000 instrument, with 90-100 millions paired reads per sample on average. Raw (FASTQ files) and processed data were deposited in GEO under accession number GSE212329.

#### DNA-seq

Genomic DNA was extracted from PGK12.1 mouse embryonic stem cells using the DNeasy blood and tissue kit (Qiagen), following the manufacturer’s instructions. The DNA-seq library for whole genome sequencing was prepared with 1µg of genomic DNA, using the Illumina TruSeq DNA Library Prep kit, following the manufacturer’s instructions. Sequencing was performed in 100pb paired-end reads using an Illumina HiSeq 2500 instrument. Raw sequencing data are available through the NIH BioProject accession number PRJNA875055.

#### SNP calling

Read pairs form the DNA-seq library were trimmed using trim_galore (https://github.com/FelixKrueger/TrimGalore) (version 0.6.3) and mapping to the mm10 genome was performed using BWA mem (Li and Durbin 2009) (version 0.7.17-r1188) with parameters: -M -k 22. Alignments were saved in BAM format and filtered using samtools view (Li et al. 2009) (version 1.9) with parameters: -q 20 -f 2. In addition, alternative hits and mitochondrial reads were removed as follows: grep -v -e ‘XA:Z:’ -e ‘SA:Z:’ -e ‘chrM’. Filtered BAM files were sorted with samtools sort and duplicates removed with picard MarkDuplicates (“Picard Toolkit.” 2019. Broad Institute, GitHub Repository. https://broadinstitute.github.io/picard/; Broad Institute) (version 2.20.8). To identify 129/PGK SNPs, bcftools (Li 2011) (version 1.9) and freebayes (Garrison and Marth 2012) (version 1.3.2) were implemented. For bcftools, the BAM file and the mm10 genome were indexed with samtools index and samtools faidx, respectively. Likelihoods were computed and variants called by first running bcftools mpileup with option -Ou, and then bcftools call with options: - mv -Ob. Variants were filtered based on quality using bcftools view with the following parameters: -i ‘%QUAL>=19’ -Ob. Variant calling with freebayes was performed with default parameters and the results were also filtered by quality with vcffilter (Garrison et al. 2022) (version 1.0.0) with option -f -f “QUAL > 19”. Variants obtained with bcftools and freebayes were filtered to only keep heterozygous SNPs using bcftools view and the following parameters: -v snps -g het -M2. Heterozygous SNPs from both programs were merged using bcftools merge and converted to BED format, to generate a list of coordinates with all possible heterozygous sites (Data S1), which was used to mask the reference genome for read mapping and minimize mapping biases. In addition, the common set of SNPs from both programs was identified using bcftools isec with option -n=2. These SNPs were filtered by quality using the values computed by freebayes as follows: vcffilter -f “QUAL > 20 & DP > 10 & SAF > 0 & SAR > 0 & RPR > 1 & RPL > 1”. Public 129 SNP reference annotations were downloaded from the Wellcome Sanger Institute website (129P2_OlaHsd.mgp.v5.snps.dbSNP142.vcf.gz), to define which of the identified alleles corresponded to the 129 genotype. Only those variant positions in common with the published reference were used to assign allele-specific reads. Finally, common SNPs overlapping low complexity regions (defined by RepeatMasker annotations) were filtered out using bedtools intersect with option -v. Thus, a total of 1,019,011 common heterozygous SNPs were identified for 129/PGK (Data S2).

#### RNA-seq data processing

Libraries were processed to remove sequencing adapters from read pairs using trim_galore. Then, to remove potential rRNA contamination, reads were mapped to a reference of rDNA annotations using STAR (Dobin et al. 2013) (version 2.7.2b) with the following parameters: -- alignIntronMax 500000 --alignMatesGapMax 500000 --alignEndsType EndToEnd -- winAnchorMultimapNmax 2000 --outFilterMultimapNmax 10000 --outFilterMismatchNmax 999 --outFilterMismatchNoverLmax 0.2 --seedMultimapNmax 20000 --outSAMtype None -- outReadsUnmapped Fastx. Filtered reads were mapped to the mm10 genome using an N-masked reference that was generated based on the merged SNP annotations of the PGK12.1 cell line (Data S1) using bedtools maskfasta (version 2.29.2). To quantify gene and single repeat element expression, read pairs were mapped to the N-masked genome using STAR to report uniquely mapped reads: --alignIntronMax 500000 --alignMatesGapMax 500000 -- alignEndsType EndToEnd --outFilterMultimapNmax 1 --outFilterMismatchNmax 999 -- outFilterMismatchNoverLmax 0.03 --outSAMattributes NH HI NM MD --outSAMmultNmax 1 --outSAMtype BAM Unsorted. On the other hand, to quantify repeat expression by family/subfamily, multi-mapping read pairs were saved by randomly asigning them to one of the possible hits using the following parameters: --alignIntronMax 500000 -- alignMatesGapMax 500000 --alignEndsType EndToEnd --winAnchorMultimapNmax 2000 -- outFilterMultimapNmax 10000 --outFilterMismatchNmax 999 --outFilterMismatchNoverLmax 0.03 --seedMultimapNmax 20000 --outSAMattributes NH HI NM MD --outSAMmultNmax 1 -- outSAMtype BAM Unsorted. BAM files with mapped reads were filtered to remove singletons with samtools view and the following options: -b -f 0x2. In addition, mitochondrial reads were filtered out using samtools view and the grep command (grep -v chrM). To assign read pairs to specific alleles, filtered BAM files and the 129/PGK SNP annotations (Data S2) were used as input for SNPsplit (Krueger and Andrews 2016) (version 0.3.4) with the following options: - -paired --no_sort. SNPsplit generated one BAM file for each allele (129 and PGK) and one for unassigned reads. These 3 BAM files were merged using samtools merge to generate a BAM file with total signal. BAM files were sorted by coordinate using picard SortSam. Total and allele-specific quantifications of gene and repeat expression were performed using featureCounts (Liao et al. 2014) (version 2.0.1) with ENSEMBL gene annotations (Cunningham et al. 2022) (v98) and RepeatMasker annotations from the UCSC genome browser (Nassar et al. 2023) (discarding simple, low complexity and exonic repeats), respectively, and with the following parameters: a) for genes, -C -p -s 2 -t exon -g gene_id; b) for single repeats, -C -p -s 2; c) for repeat families/subfamilies, -C -p -s 2 -M. Repeat coordinates were specified as repeat IDs in the SAF file used by featureCounts. Family/subfamily quantification was obtained by summing counts of all repeats according to their classification. Read counts of single repeats were used to compute Transcripts Per Million (TPM) values and obtain their expression fold change upon DOX treatment. Signal profiles were generated for each library and by strand using bamCoverage (Ramirez et al. 2016) (version 3.1.3), a scaling factor (100000000/library size) and the following parameters: --binSize 1 --normalizeUsing CPM --filterRNAstrand [forward or reverse] -- effectiveGenomeSize 2467481108 --outFileFormat bigwig --scaleFactor [computed for each library].

#### Normalization by repeat family and subfamily

Read count tables of repeat families and subfamilies were processed to add up reads of repeat elements grouped by family or subfamily and obtain their total quantification. The two tables generated were independently joined with the read count table of genes to perform normalization. The R package DESeq2 (Love et al. 2014) (version 1.26.0) was used to calculate normalized counts for all libraries by clone. Normalized counts were extracted from the DESeqDataSet using the function counts with the option normalized=TRUE, and only the normalized counts of repeat families and subfamilies were used for further analyses.

#### Differential gene and repeat expression

Contrast analyses of libraries with and without DOX treatment by clone were performed using DESeq2. To generate one input table for DESeq2, read count tables of genes and single repeats were joined. Genes and repeats with less than 10 total counts across libraries were discarded for comparison. DE analysis was performed using the DESeq function and computed values extracted using the results function. To calculate corrected p values, genes and repeats discarded by independent filtering or considered as outliers (NA values in p value and adjusted p value columns) were removed from the results table. Then, the full column of adjusted p values was deleted and new p values were estimated using the z-scores with the function fdrtool setting statistic = “normal”. Adjusted p values were then computed using the Benjamini-Hochberg method with the function p.adjust. Results of genes were extracted and visualized using volcano plots generated with the function EnhancedVolcano (https://github.com/kevinblighe/EnhancedVolcano) and setting the option pCutoff = 0.01, to add a line at the adjusted p value threshold used to consider genes as differentially expressed.

#### Metagene plot signal

To generate repeat expression profiles, coordinates of differentially expressed L1MdTfs by strand and clone were converted to BED format. In the case of the repressor clone, only the downregulated repeats were considered, whereas for the activator clones, only the upregulated repeats were considered. deepTools (version 3.4.3) was used to compute average values and generate two signal plots by clone, one in sense and one in antisense orientation. First, average values by strand were calculated using computeMatrix scale-regions with options: -m 6000 -b 2000 -a 2000 --missingDataAsZero. For sense transcription, signal profiles (bigWig files) corresponding to the strand of the annotated L1MdTfs were used as computeMatrix input, while signal profiles of the opposite strand of the annotated L1MdTfs were used to calculate the antisense signal. Then, values obtained for each of the BED files containing DE L1MdTfs in the forward and reverse strand were joined using computeMatrixOperations rbind. Finally, plots were generated using plotProfile with options: - -averageType mean --perGroup.

#### Clustering of genes and repeats

Fold changes of DE genes and L1MdTf repeats were used to compute z-scores, to perform K-means clustering using the kmeans function in R (R Core Team (2023)) (version 3.6.1). The number of clusters was selected based on visual assessment (heat maps) of the generated clusters. The joint table of raw read counts of genes and single repeats of all clones was filtered to remove features with less than 10 total reads across libraries. A regularized log transformation was applied to this filtered table using the DESeq2 rlog function with option: blind=FALSE. Normalized read counts of DE genes and L1MdTf repeats were extracted and used to generate gene expression heat maps based on the order obtained by clustering their corresponding z-scores.

#### Distance analyses

Genomic distances between L1MdTf repeats and all genes and non-L1 repeats, which were included in the DE analyses, were calculated using bedtools (version 2.29.0) function closest with the following parameters: -d t “first”. BED files used for these analyses contained the coordinates of full gene/repeat annotations. Genes and non-L1 repeats were classified according to the DE results, significance of the observed differences in the distance distribution of classes was assessed using Wilcoxon-test (wilcox.test function in R). To test the dependence between distance and gene/repeat classification, Chi-squared tests were performed using the chisq.test function in R with option: simulate.p.value = TRUE. Residuals from this test were plotted using the corrplot package (https://github.com/taiyun/corrplot) (version 0.88).

#### Chromatin state analysis

The ChromHMM model of ESCs used for analyses was downloaded from: https://github.com/guifengwei/ChromHMM_mESC_mm10. Fold enrichment of DE genes and non-L1 repeats in the 12 annotated states was tested using ChromHMM OverlapEnrichment (Ernst and Kellis 2017) (version 1.23).

#### Gene ontology enrichment analysis

Enrichment analyses of GO terms, KEGG pathways and Uniprot tissue expression were performed with the Database for Annotation, Visualization and Integrated Discovery – DAVID (Sherman et al. 2022) (version 6.8) by providing the ENSEMBL IDs of DE genes in the activator system and using the whole mouse genome as background. Results tables were downloaded from the online tool and adjusted p values, fold enrichment values and number of genes by term were used for visualization.

#### Allelic expression analyses

Allele-specific repeat and gene read counts of 129 and PGK obtained with featureCounts were used to calculate allelic ratios and test for differences from the expected biallelic expression with DESeq2. For each clone, a DDS object was generated using its read count table and a design matrix specifying the DOX treatment, replicate information and the allele. The design formula for contrasts was set as follows: ∼treatment + treatment:replicate + treatment:allele. Genes and repeats with less than 10 reads across samples were removed from the DDS object. Then, size factors were set to 1 and significance testing performed with the DESeq function. Results tables were extracted with the results function and the following options: alpha = 0.01, filterFun = ihw (Ignatiadis et al. 2016). The obtained adjusted p values represent genes or repeats with significant bias from biallelic expression (allelic score = 0.5). PGK/129 ratios were calculated from the log2 fold changes in the results table and allelic scores were then calculated as: (PGK/129) / (1 + PGK/129). Genes and repeats with adjusted p values lower than 0.01 were considered as features with skewed allelic-expression.

#### Data visualization

Results from bioinformatics analyses were plotted using R (version 3.6.1) and the ggplot2 package (https://ggplot2.tidyverse.org/) (version 3.1.1).

## Acknowledgments

We are grateful to members of the Heard lab, where this project was initiated and Y.A.P.-R., A.B. and E.M. worked as paid postdocs of the lab and A.-V.G. as researcher; Rafael Galupa (CBI, Toulouse) for critical reading of the manuscript; Elphège Nora (UCSF) for sharing the sgRNAs and donor plasmids for *Rosa26* and *Tigre*; Michel Wassef (Institut Curie) for cloning of the pEN366-dCas9-VPR inducible construct; and Aurélie Tessandier (Institut Curie) for help with the initial mapping of the L1 sgRNAs. High-throughput sequencing was performed by the ICGex NGS platform of Institut Curie supported by the grants ANR-10-EQPX-03 (Equipex) and ANR-10-INBS-09-08 (France Génomique Consortium) from the Agence Nationale de la Recherche (“Investissements d’Avenir’’ program), by the ITMO-Cancer Aviesan (Plan Cancer III) and by the SiRIC-Curie program (SiRIC Grant INCa-DGOS-465 and INCa-DGOS-Inserm_12554). Data management, quality control and primary analysis were performed by the Bioinformatics platform of Institut Curie. We also thank the Bioimaging facility at iMM-JLA for their services and assistance.

This project has received funding from the European Union’s Horizon 2020 research and innovation programme under the Marie Skłodowska-Curie grant agreement No 882771 to Y.A.P-R; from national Portuguese funds through FCT-Fundação para a Ciência e a Tecnologia, CEECIND/02085/2018 and 2022.02195.PTDC to A.-V.G; from an ERC Advanced Investigator award, ERC-ADG-2014 671027 to E.H; and from NIH R01AG078959 to A.R.M.

## Supplemental figures

**Figure S1.**
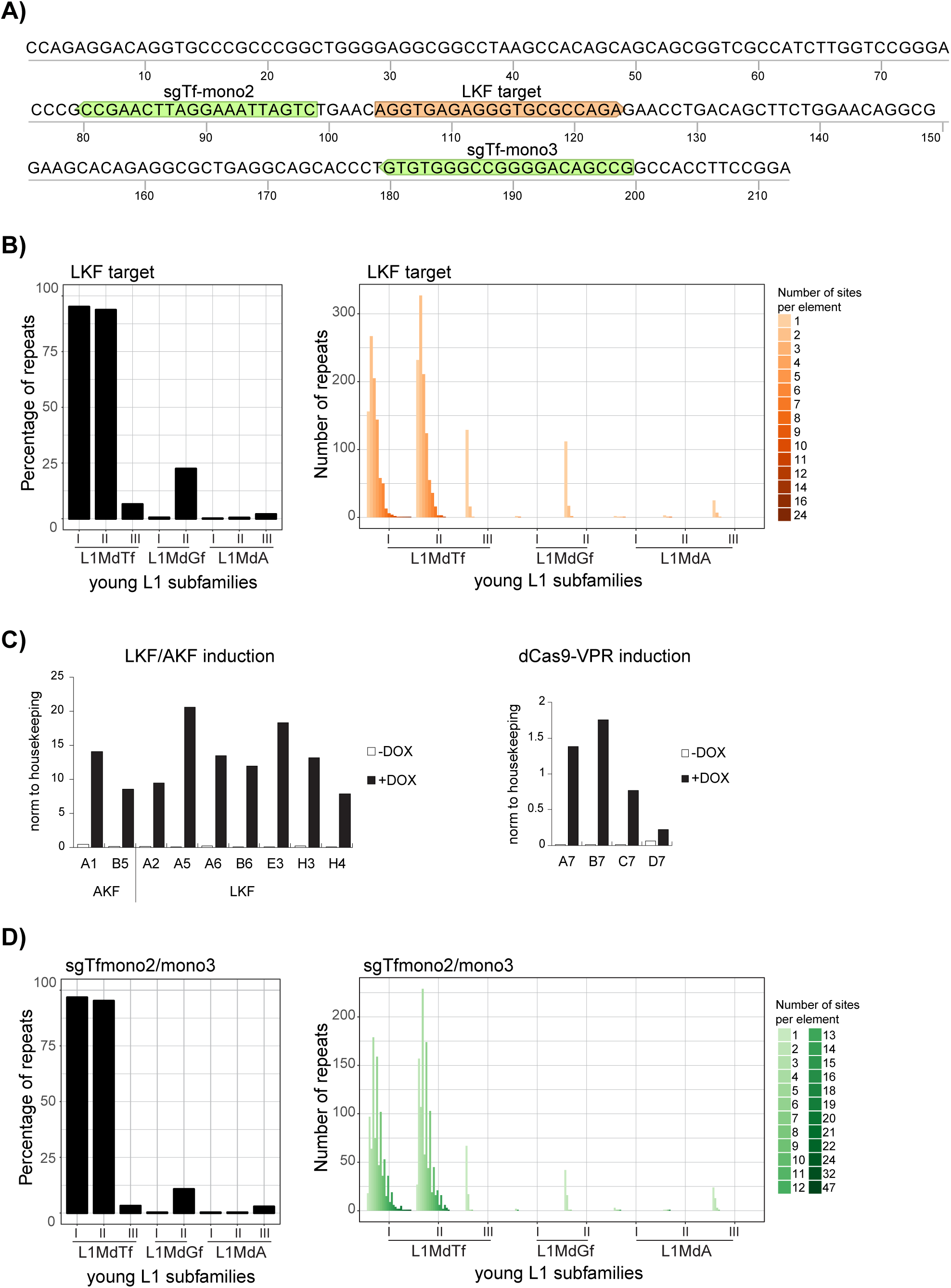
Identity and genomic representation of L1MdTf-repressor and activator target sequences. A) Positions of LKF and sgRNAs (Tf-mono2 and Tf-mono3) target sequences within the 212 bp L1MdTf monomer consensus sequence. B) Estimation of the percentage of young L1Md subfamilies bound by LKF repressor (left). Predicted number of elements from young L1Md subfamilies bound by LKF repressor according to the number of sites per element (right). C) Following puromycin selection, colony picking and genotyping, 9 clones were analysed upon DOX treatment (48h) by qPCR for expression of the LKF/AKF transgenes (left) and 4 clones for the expression of the dCas9-VPR transgene (right). D) Estimation of the percentage of young L1Md subfamilies targeted by Tfmono2/3 guide RNAs (left). Number of elements from young L1Md subfamilies targeted by Tfmono2/3 according to the number of sites per element (right).

**Figure S2.**
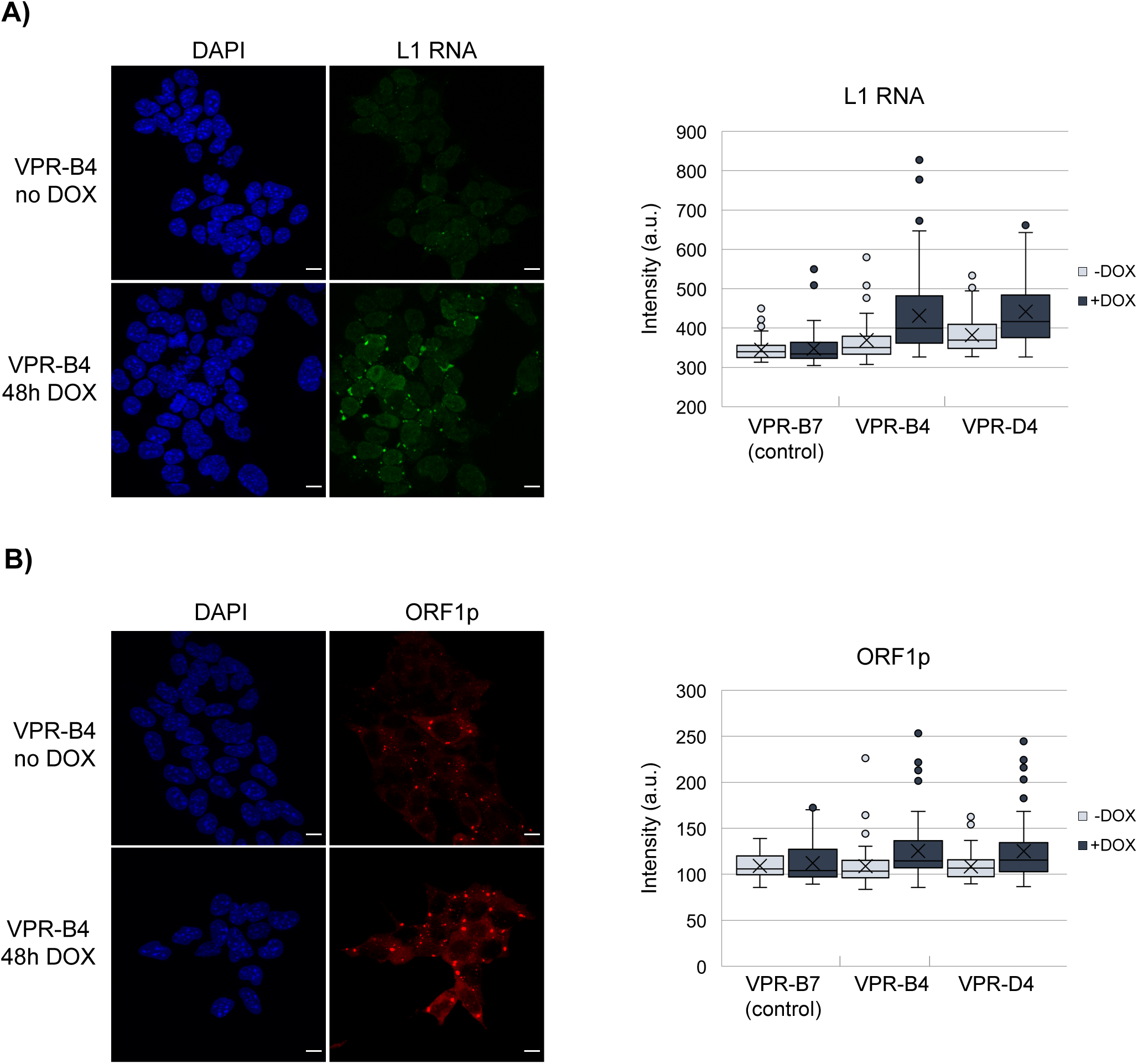
Visualization of L1 RNA and ORF1p expression in control and activator clones. A) Representative images of L1 RNA FISH in activator clone VPR-B4 in the presence or absence of DOX for 48h (left) (scale bar, 10 μm). Quantification of L1 RNA signal, detected with full-length L1 probe (TNC7), in control and activator clones based on the average of the mean grey values per cell and their standard deviation for each clone/condition (n=80-200 cells) (right). B) Representative images of ORF1 immunofluorescence in activator clone VPR-B4 in the presence or absence of DOX for 48h (left) (scale bar, 10 μm). Quantification of ORF1 signal in control and activator clones based on the average of the mean grey values per cell and their standard deviation for each clone/condition (n=50-90 cells) (right).

**Figure S3.**
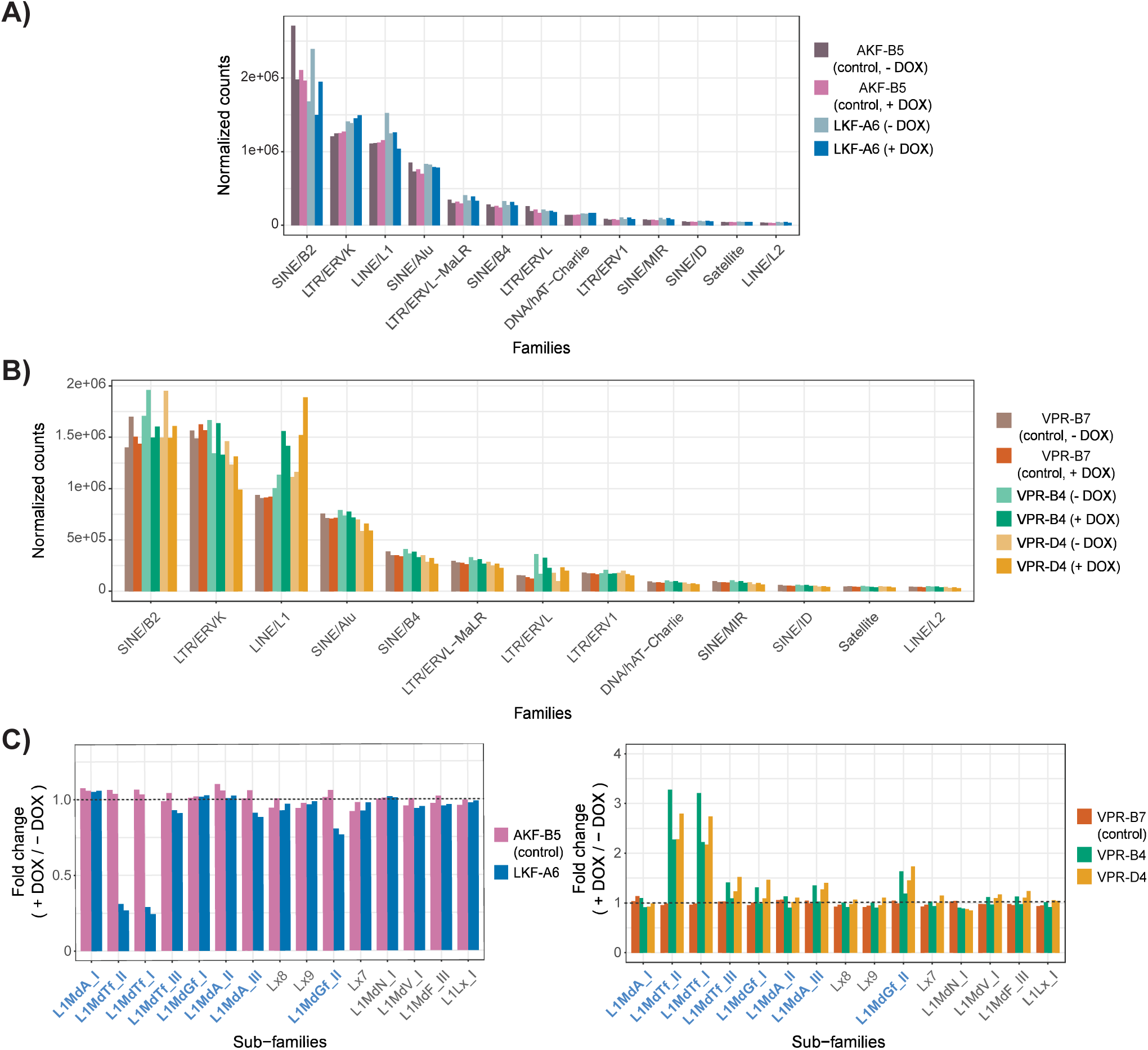
Normalized RNA-seq expression by family and subfamily of repeats. A) Bar plot showing normalized read counts (y-axis) per repeat family in repressor (LKF-A6) and control (AKF-B5) clones with and without DOX treatment. B) Bar plot showing normalized read counts (y-axis) per repeat family in activator (VPR-B4, VPR-D4) and control (VPR-B7) clones with and without DOX treatment. C) Bar plots showing fold change of normalized read counts after DOX treatment (y-axis) per L1 subfamily in repressor (left) and activator (right) clones with their respective controls. Dash lines mark values equal to 1.

**Figure S4.**
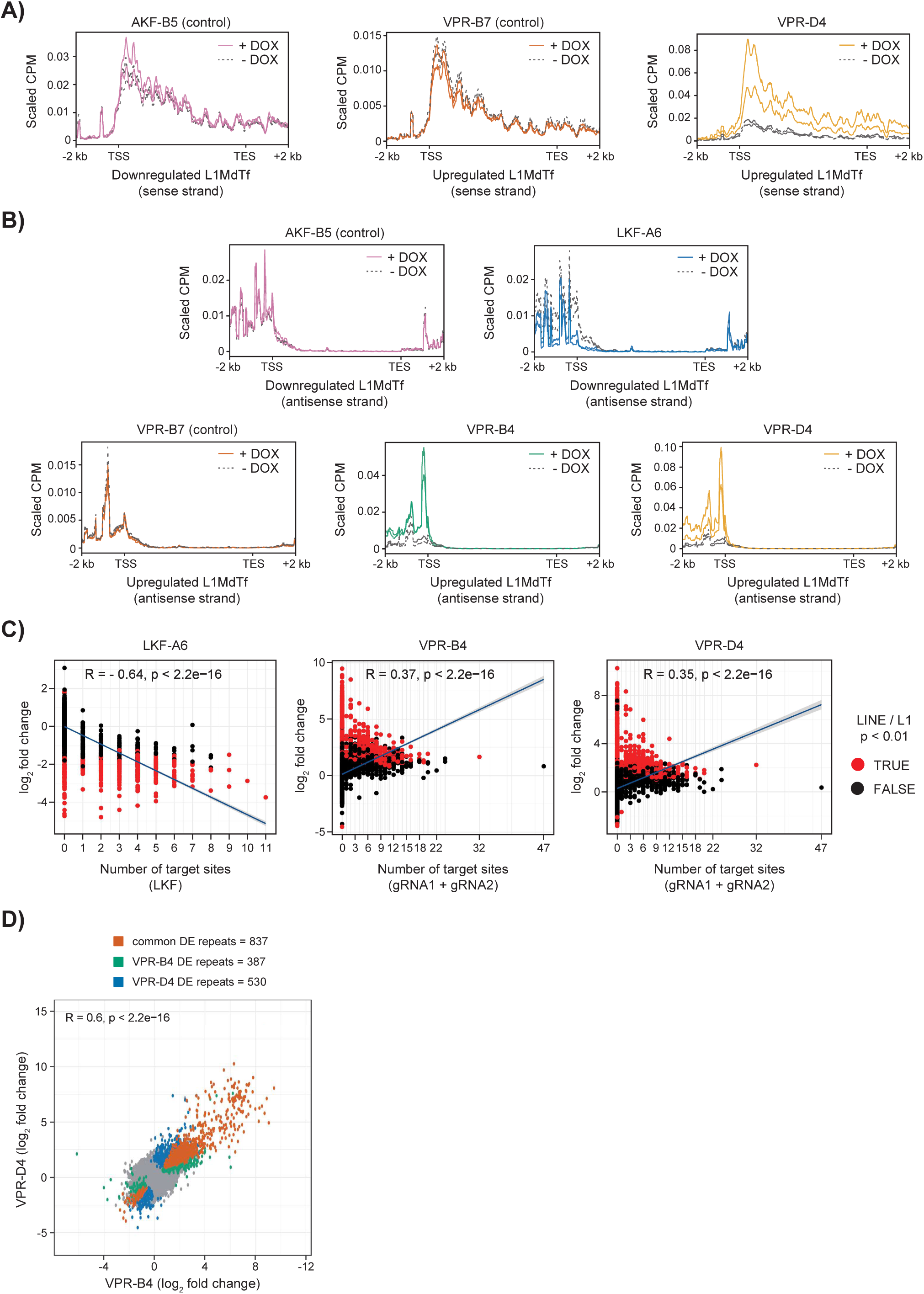
Analysis of L1MdTf repeat expression after DOX treatment. A) Metagene plot of normalized RNA-seq signal (shown for AKF-B5, VPR-B7 and VPR-D4 clones, replicates A & B) on L1MdTf repeats showing downregulation in LKF-A6 (left) or upregulation in the activator clones (middle, right) upon DOX treatment. The x-axis is scaled to represent full annotations and the 2 kb upstream and downstream regions. B) Metagene plot of normalized RNA-seq signal (shown for all clones sequenced, replicates A & B) on the antisense strand of L1MdTf repeats showing downregulation in the LKF-A6 clone (top) or upregulation in the activator clones (bottom) upon DOX treatment. The x-axis is scaled to represent full annotations and the 2 kb upstream and downstream regions. C) Correlation between expression fold change and number of LKF (left) or gRNA (middle, right) target sites for L1 repeats with (red) and without (black) significant changes in expression in repressor (left) and activator (middle, right) clones upon DOX treatment. Pearson correlation values and their significance are displayed. D) Comparison of expression fold changes of repeats in the VPR-B4 and VPR-D4 clones after DOX treatment. Grey dots represent repeats with no significant changes in expression and colored dots represent repeats identified as DERs in both clones or in one of the clones only. Pearson correlation coefficient and significance are shown.

**Figure S5.**
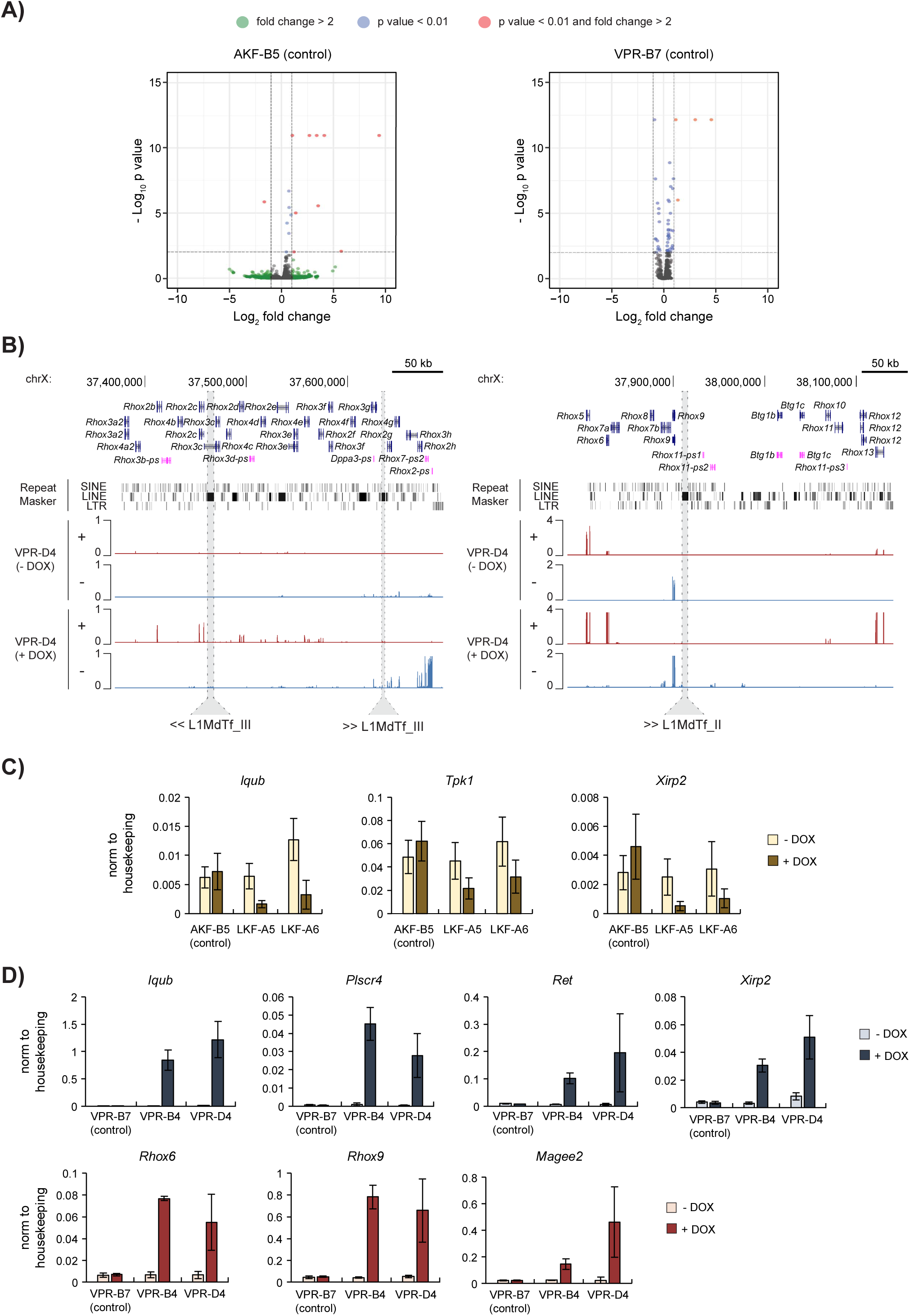
DE analysis of genes in control clones and examples of selected and validated DEGs. A) Volcano plots showing gene expression fold changes (x-axis) and their significance (y-axis) in the repressor AKF-B5 (left) and activator VPR-B7 (right) control clones. B) Examples of upregulated genes in the *Rhox* cluster on the X chromosome following activation of L1MdTf repeats. RNA-seq profiles by strand are shown in red (+ strand) and blue (- strand) for the DOX-treated and untreated samples. L1MdTf repeats located in the cluster and their orientation are highlighted. C) qPCR results for selected genes showing specific downregulation in repressor clones, following DOX treatment for 48h. Levels are normalized relative to housekeeping gene expression. Error bars represent the calculated standard deviation considering 6 biological replicates. D) qPCR results for selected genes (top, autosomal genes; bottom, X-linked genes) showing specific upregulation in the activator clones, following DOX treatment for 48h. Levels are normalized relative to housekeeping gene expression and error bars represent the calculated standard deviation considering 3 biological replicates.

**Figure S6.**
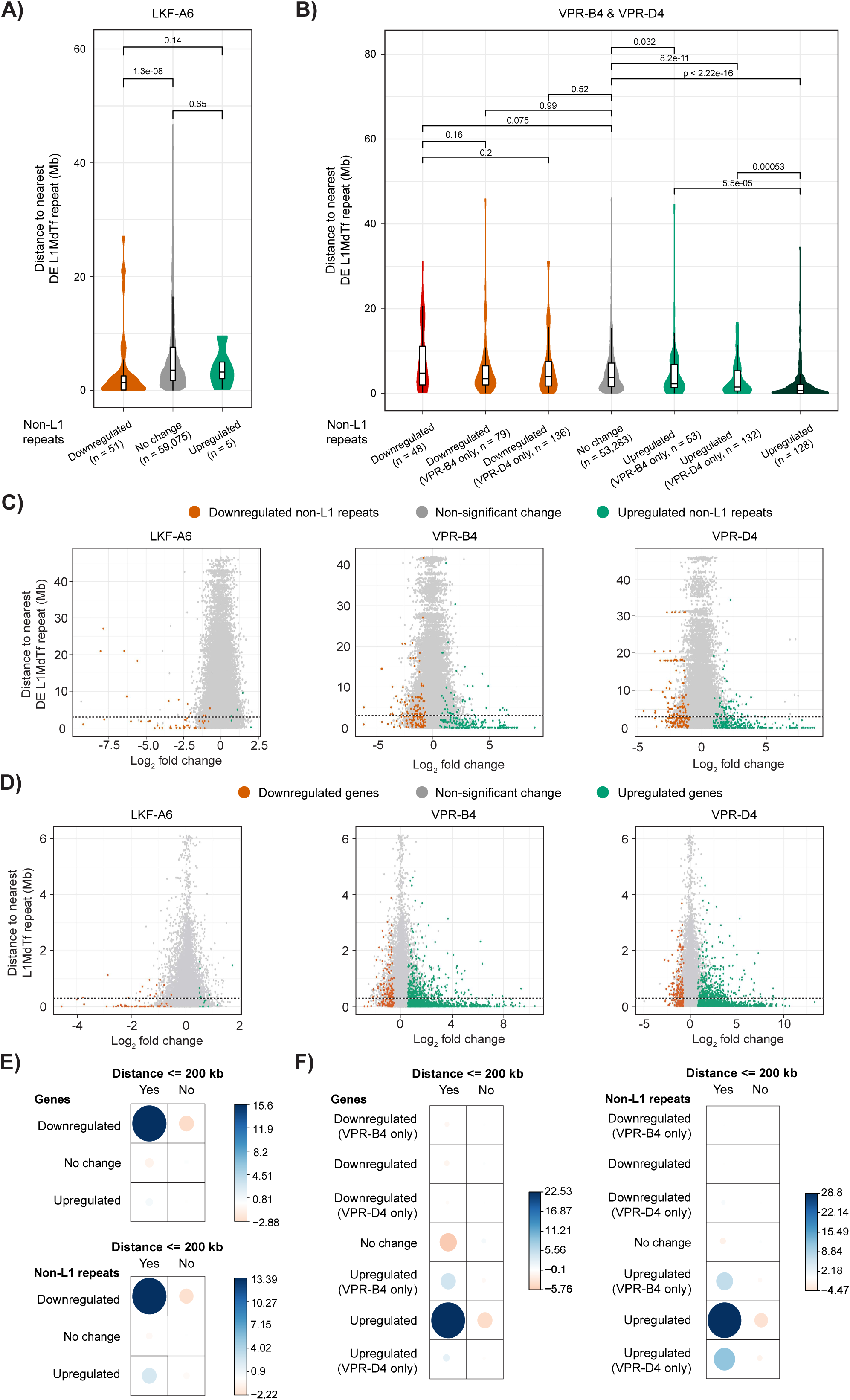
Distance analyses between non-L1 DERs and DE L1MdTf repeats. A) Distribution of distances between DE L1MdTf repeats and non-L1 repeats classified according with DE analysis results in the LKF-A6 clone. Significance values of the differences between distributions are displayed above violin plots and were calculated using Wilcoxon rank-sum tests. B) Distribution of distances between DE L1MdTf repeats and non-L1 repeats classified according with DE analysis results in the activator clones. Significance values as described for panel A. C) Comparison between expression fold change upon L1MdTf repression (LKF-A6) or activation (VPR-B4, VPR-D4) and distance to nearest DE L1MdTf repeat for all non-L1 repeats tested for DE. Dashed lines indicate a distance of 3 Mb. D) Comparison between expression fold change upon L1MdTf repression or activation and distance to nearest annotated L1MdTf repeat for all genes tested for DE. Dashed lines indicate a distance of 260 kb. E) Pearson residuals from Chi-square tests representing the relevance of attributes (i.e. classification based on expression and distance) to the total Chi-square score for genes (top) and non-L1 repeats (bottom) in the repressor clone. Bigger circles depict higher contribution of table cells to the score. Colors indicate positive and negative association between rows and columns. F) Pearson residuals from Chi-square tests as described on panel E for genes and non-L1 repeats in the activator clones.

**Figure S7.**
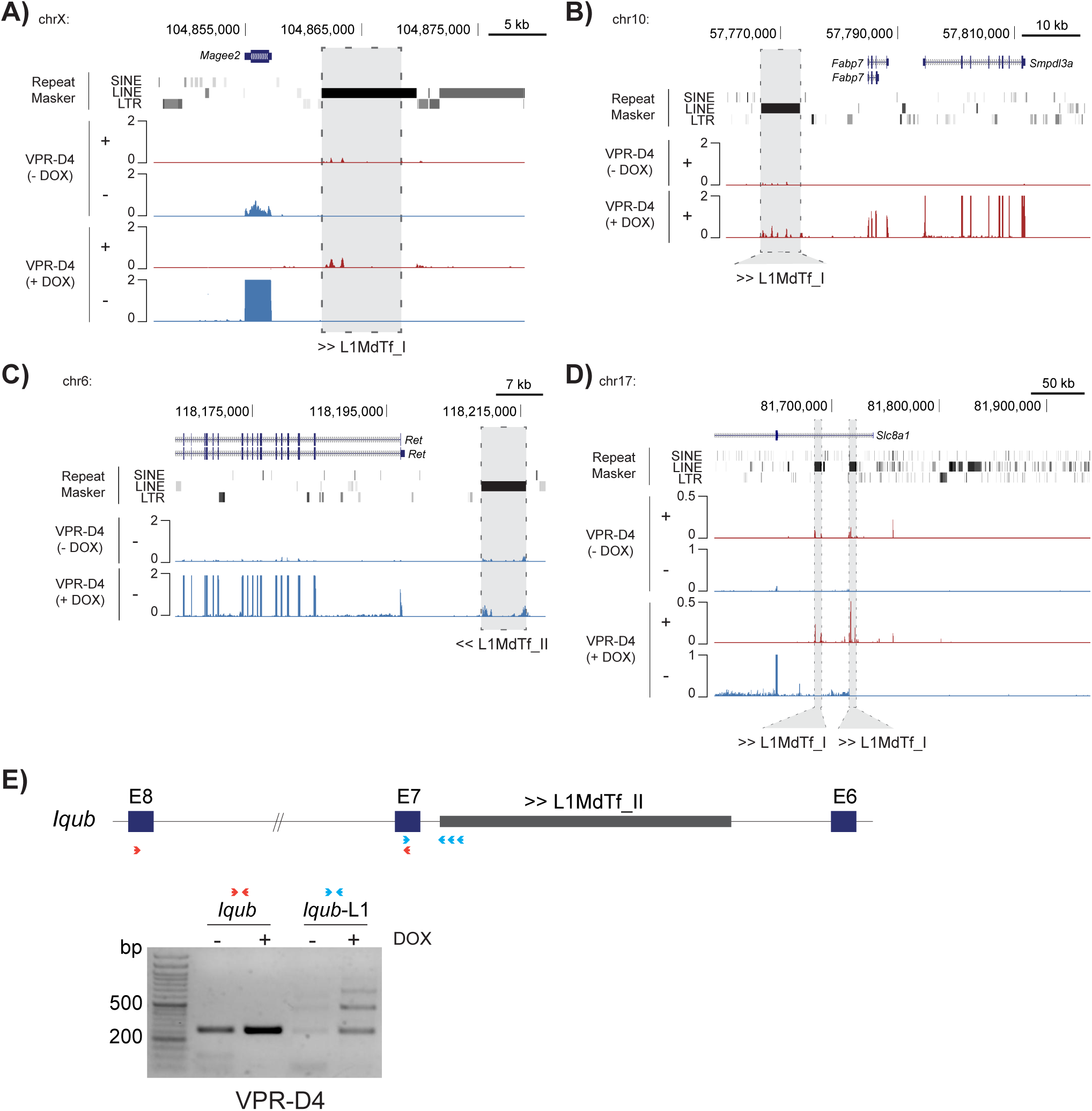
Genome browser screenshots for selected genes proximal to misregulated L1MdTf repeats. A) *Magee2* vicinity showing increased expression upon L1MdTf activation. RNA-seq profiles by strand are shown in red (+ strand) and blue (- strand) for DOX-treated and untreated activator samples. An adjacent L1MdTf repeat with reciprocal expression change, and its orientation, is highlighted. B) *Fabp7*/*Smpdl3a* and C) *Ret* vicinities showing increased expression upon L1MdTf activation as on panel A. D) *Slc8a1* vicinity showing an example of a gene with increased expression upon L1MdTf activation, which contains intronic L1MdTfs with reciprocal changes and located in inverse orientation with respect to *Slc8a1*. E) Semi-quantitative RT-PCR analysis of the *Iqub* locus in the VPR-D4 clone using two different sets of primers. The red primer pair overlaps *Iqub* exonic regions and yields a PCR amplicon of 263 bp, while the blue primer pair overlap an *Iqub* intron and the repetitive monomers within the 5’UTR of the intronic L1 element, yielding products of 254, 466 and 678 bp.

**Figure S8.**
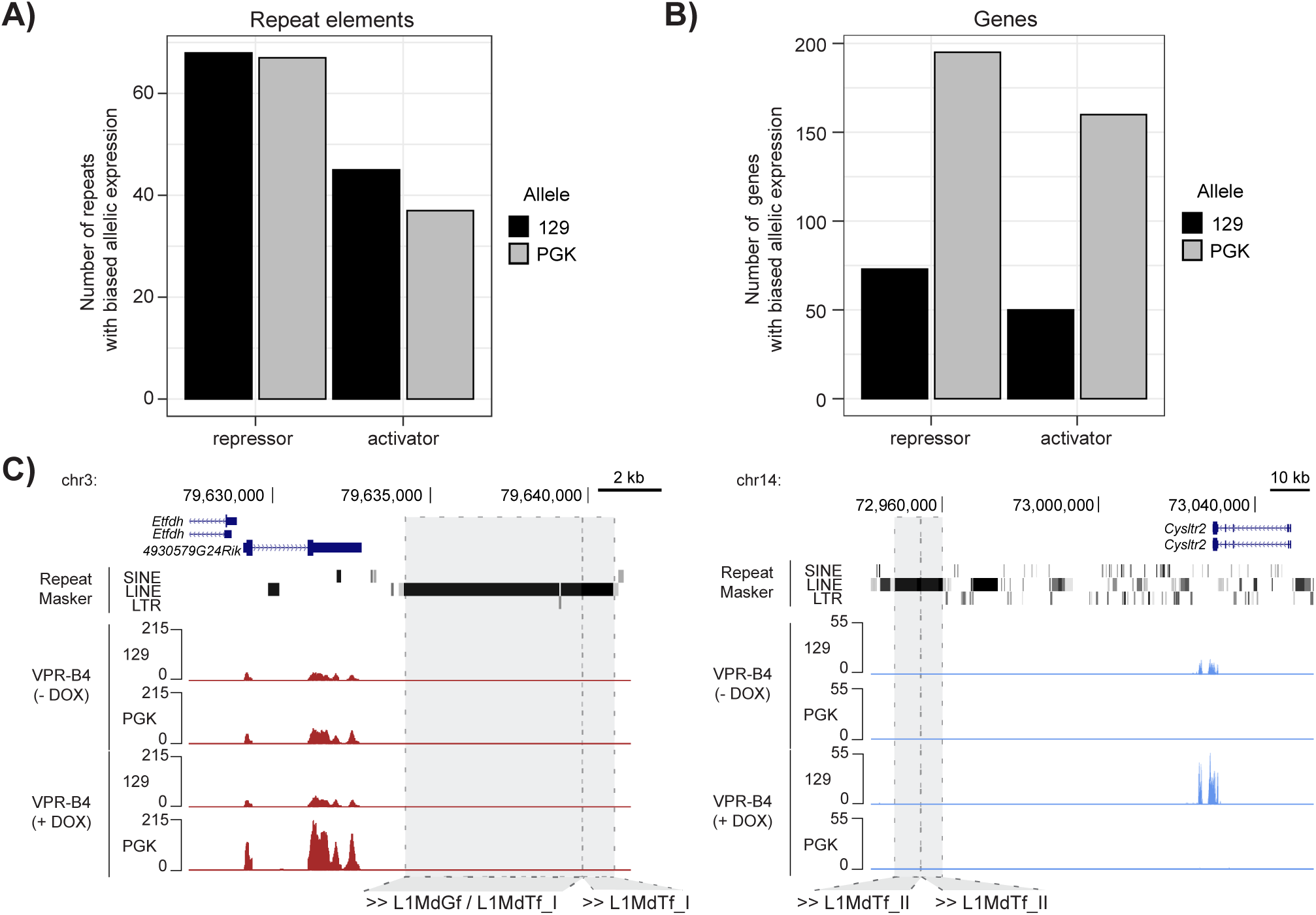
Identification of repeats and genes with skewed allelic expression. A) Repeat elements with skewed allelic expression in L1MdTf repressing and activating conditions. Allele classification (129 or PGK) shows the allele from which elements are preferentially expressed. B) Genes with skewed allelic expression as described in A. C) Genome browser screenshots for the *4930579G24Rik* and *Cysltr2* genes, which show skewed expression following dCas9-VPR-mediated activation of L1MdTf repeats.

**Figure S9.**
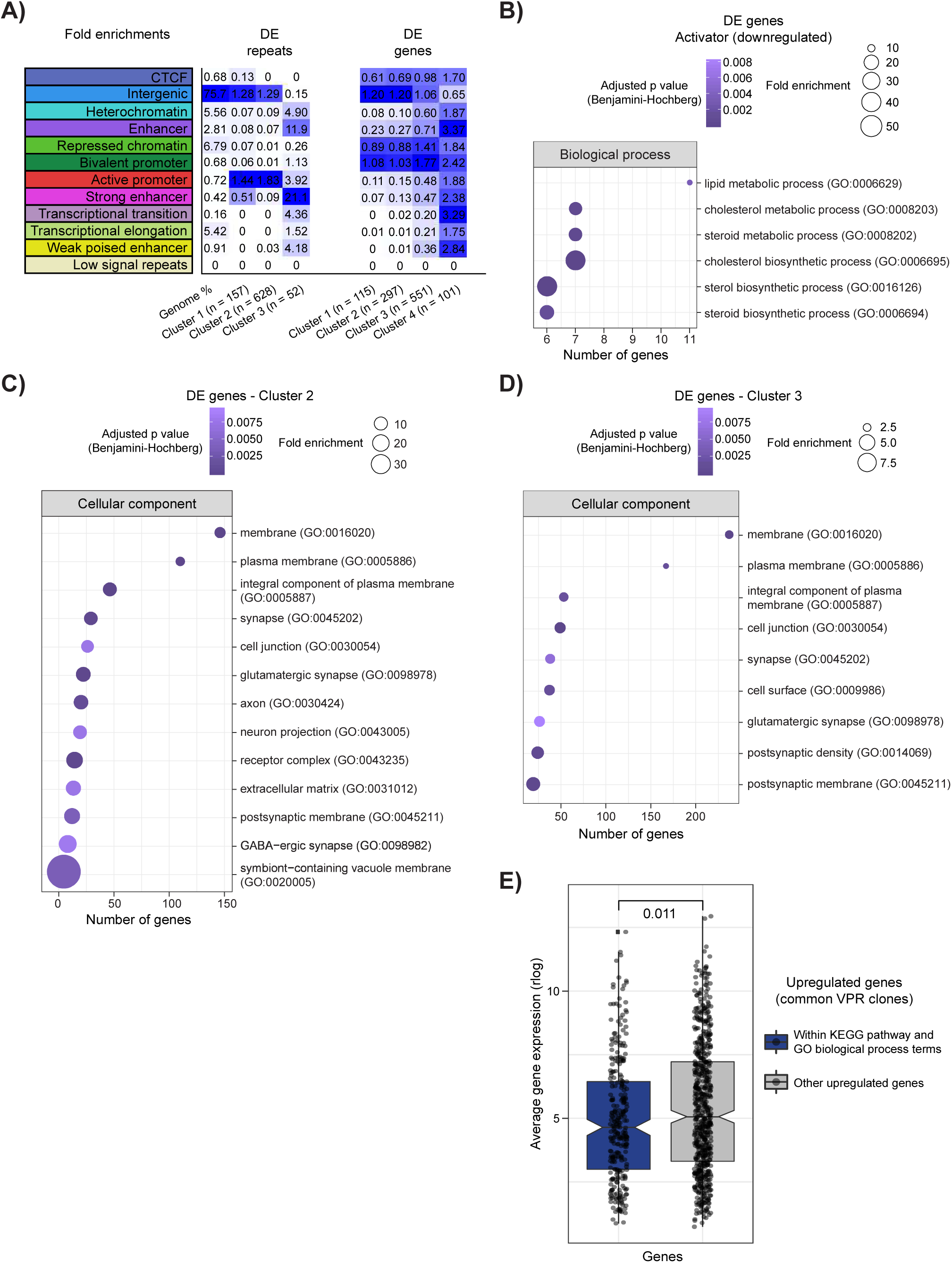
Chromatin state and GO enrichment analyses for DEG and DER clusters. A) Heat map showing fold enrichment for each chromatin state of DE repeat and gene clusters shown on Fig.3E and Fig.4E. Increased color darkness represents increased enrichment. B) GO enrichment analysis of biological process terms for the common downregulated genes in activator clones. The number of genes by term is shown on the x-axis, circle size indicates fold enrichment and darker colors indicate higher significance. C) GO enrichment analysis of cellular component terms for the genes in cluster 2. D) GO enrichment analysis of cellular component terms for the genes in cluster 3. For C and D, plot annotations are as described for panel B. E) Box plot showing the distribution of average expression of genes that are part of the enriched GO biological process terms or KEGG pathways (shown on Fig.6C) and the rest of genes classified as upregulated in the activator clones. P value testing the significance of the observed difference in distributions is displayed above the boxes.

## Supplemental tables

**Table S1**- List of predicted LKF repressor target sites in the mouse genome (mm10). Coordinates of the target sites are included, as well as, the information of the overlapping repeat element.

**Table S2**- Joint list of predicted target sites of the two L1 sgRNAs used for the VPR clones in the mouse genome (mm10). Coordinates of the target sites are included, as well as, the information of the overlapping repeat element.

**Table S3**- DESeq2 normalized read counts per repeat family in all RNA-seq libraries. Biological replicates are indicated with the suffixes “_A” and “_B”.

**Table S4**- DESeq2 normalized read counts per repeat subfamily including family classification in all RNA-seq libraries. Biological replicates are indicated with the suffixes “_A” and “_B”.

**Table S5**- Differential expression analysis results of genes and repeats for LKF-A6 samples with and without DOX treatment. The results include normalized read counts (baseMean), fold changes (log2FoldChange) and adjusted p values (padj). IDs correspond to ENSEMBL ids for genes and genomic coordinates for repeats. Name and family/subfamily are indicated for genes and repeats, respectively.

**Table S6**- Differential expression analysis results of genes and repeats for VPR-B4 samples with and without DOX treatment. The results include normalized read counts (baseMean), fold changes (log2FoldChange) and adjusted p values (padj). IDs correspond to ENSEMBL ids for genes and genomic coordinates for repeats. Name and family/subfamily are indicated for genes and repeats, respectively.

**Table S7**- Differential expression analysis results of genes and repeats for VPR-D4 samples with and without DOX treatment. The results include normalized read counts (baseMean), fold changes (log2FoldChange) and adjusted p values (padj). IDs correspond to ENSEMBL ids for genes and genomic coordinates for repeats. Name and family/subfamily are indicated for genes and repeats, respectively.

**Table S8**- List of the 206 DERs with reciprocal changes in repressor (LKF-A6) and activator (VPR-B4, VPR-D4) clones. Coordinates, classification and the assigned cluster number (Fig.3D) are included for each repeat.

**Table S9**- List of the 837 common DERs in the activator (VPR-B4, VPR-D4) clones. Coordinates, classification and the assigned cluster number (Fig.3E) are included for each repeat.

**Table S10**- Differential expression analysis results of genes and repeats for AKF-B5 samples with and without DOX treatment. The results include normalized read counts (baseMean), fold changes (log2FoldChange) and adjusted p values (padj). IDs correspond to ENSEMBL ids for genes and genomic coordinates for repeats. Name and family/subfamily are indicated for genes and repeats, respectively.

**Table S11**- Differential expression analysis results of genes and repeats for VPR-B7 samples with and without DOX treatment. The results include normalized read counts (baseMean), fold changes (log2FoldChange) and adjusted p values (padj). IDs correspond to ENSEMBL ids for genes and genomic coordinates for repeats. Name and family/subfamily are indicated for genes and repeats, respectively.

**Table S12**- List of the 31 genes identified as differentially expressed in repressor (LKF-A6) and activator (VPR-B4, VPR-D4) clones. Coordinates, gene name and the assigned cluster number (Fig.4C) are included for each gene.

**Table S13**- List of the 1,024 common DEGs in the activator (VPR-B4, VPR-D4) clones. Coordinates, gene name and the assigned cluster number (Fig.4E) are included for each gene.

**Table S14**- List of annotated genes and repeats (excluding LINE/L1) tested for differential expression in the LKF-A6 samples showing their distance to a DE L1MdTf repeat (dist_DE_L1MdTf) and all annotated L1MdTfs (dist_L1MdTf). Overlapping elements are indicated with zero.

**Table S15**- List of annotated genes and repeats (excluding LINE/L1) tested for differential expression in the VPR-B4 samples showing their distance to a DE L1MdTf repeat (dist_DE_L1MdTf) and all annotated L1MdTfs (dist_L1MdTf). Overlapping elements are indicated with zero.

**Table S16**- List of annotated genes and repeats (excluding LINE/L1) tested for differential expression in the VPR-D4 samples showing their distance to a DE L1MdTf repeat (dist_DE_L1MdTf) and all annotated L1MdTfs (dist_L1MdTf). Overlapping elements are indicated with zero.

**Table S17**- List of annotated genes and repeats (excluding LINE/L1) tested for differential expression in both activator samples (VPR-B4, VPR-D4) showing their distance to a DE L1MdTf repeat (dist_DE_L1MdTf) and all annotated L1MdTfs (dist_L1MdTf). Overlapping elements are indicated with zero.

**Table S18**- GO enrichment analysis results of biological process terms for the common downregulated genes in the activator clones.

**Table S19**- Enrichment analysis results of GO biological process terms, Uniprot tissue expression and KEGG pathways for the common upregulated genes in the activator clones.

**Table S20**- GO enrichment analysis results of cellular component terms for the common upregulated genes in cluster 2 of the activator clones.

**Table S21**- GO enrichment analysis results of cellular component terms for the common upregulated genes in cluster 3 of the activator clones.

**Table S22**- List of guide RNA sequences and genotyping primers.

**Table S23**- List of qPCR and RT-PCR primers.

## Supplemental Data

**Data S1-** Coordinate list of all heterozygous SNPs identified with bcftools and freebayes in BED format.

**Data S2-** Common heterozygous SNPs identified with bcftools and freebayes formatted to be used as input of SNPsplit.

